# Identification of Genetic and Genomic Influences on Progressive Ethanol Consumption in Diversity Outbred Mice

**DOI:** 10.1101/2023.09.15.554349

**Authors:** ML Smith, Z Sergi, KM Mignogna, NE Rodriguez, Z Tatom, L MacLeod, KB Choi, V Philip, MF Miles

**Author notes:** **Corresponding author:** Michael Miles, MD, PhD **E-mail:**. **Abbreviations:** AUD = alcohol use disorder; Chr = chromosome; DE = differentially expressed; DO = Diversity Outbred; EtOH = ethanol; eQTL = expression quantitative trait loci; genome construction by RNAseq = GBRS; GO = Gene Ontology; IEA = intermittent access ethanol; KDA = key driver analysis; MEs = module eigengenes; PFC = prefrontal cortex; SNP = single nucleotide polymorphism; TPM = transcripts per million; VST = variance stabilizing transformation; WGCNA = Weight Gene Correlated Network Analysis.

## Abstract

Genetic factors play a significant role in the risk for development of alcohol use disorder (AUD). Using 3-bottle choice intermittent access ethanol (IEA), we have employed the Diversity Outbred (DO) mouse panel as a model of alcohol use disorder in a genetically diverse population. Through use of gene expression network analysis techniques, in combination with expression quantitative trait loci (eQTL) mapping, we have completed an extensive analysis of the influence of genetic background on gene expression changes in the prefrontal cortex (PFC). This approach revealed that, in DO mice, genes whose expression was significantly disrupted by intermittent ethanol in the PFC also tended to be those whose expression correlated to intake. This finding is in contrast to previous studies of both mice and nonhuman primates. Importantly, these analyses identified genes involved in myelination in the PFC as significantly disrupted by IEA, correlated to ethanol intake, and having significant eQTLs. Genes that code for canonical components of the myelin sheath, such as *Mbp*, also emerged as key drivers of the gene expression response to intermittent ethanol drinking. Several regulators of myelination were also key drivers of gene expression, and had significant QTLs, indicating that genetic background may play an important role in regulation of brain myelination. These findings underscore the importance of disruption of normal myelination in the PFC in response to prolonged ethanol exposure, that genetic variation plays an important role in this response, and that this interaction between genetics and myelin disruption in the presence of ethanol may underlie previously observed behavioral changes under intermittent access ethanol drinking such as escalation of consumption.

## Introduction

Alcohol use disorder (AUD) is a clinically significant condition characterized by an inability to stop or control one’s alcohol use in spite of negative consequences [1]. While human studies have identified a number of environmental factors that increase risk for AUD [2], genetics also play a significant role with 40-60% of AUD risk attributable to heritable factors [3, 4]. Genome-wide association studies in human subjects have identified a growing number of genetic variants associated with alcohol consumption and AUD, the most replicable of which are in alcohol dehydrogenase (ADH) and aldehyde dehydrogenase (ALDH) genes [5, 6], and, more recently, in genes such as *AUTS2, KLB,* and *CADM2*. However, these studies have generally found only small effect sizes for AUD-associated variants in ADH and ALDH genes [5], and other potential candidates genes identified by individual GWAS have been difficult to replicate [7, 8]. There are a number of factors that may be the cause, such as that alcohol use disorder is a heterogeneous condition with varying symptomatology, including the length-of-time and severity of problematic alcohol use [9].

Animal models provide a means of controlling for certain factors that cannot be in human patients. Mouse models, in particular, have been useful for identifying physiological mechanisms of aspects of AUD such as analgesia [10], anxiolysis [11], and withdrawal [12]. Mice have also been used to explore the transcriptomic changes that occur in the brain with acute [13] and chronic ethanol exposure [14–16]. However, traditional inbred mouse lines or genetic panels may not be the best models for genetic association studies due to their homozygosity and relative lack of genetic variation. To this end, the Diversity Outbred (DO) mouse panel was created using 5 common inbred laboratory mouse strains (A/J, C57BL/6J, 129S1/SvlmJ, NOD/ShiLtJ, NZO/HILtJ) and 3 wild-derived inbred strains (CAST/EiJ, PWK/PhJ, WSB/EiJ). After multiple generations of randomized breeding, the result was a mouse population with greater genetic diversity than traditional inbred mouse strains and a high degree of heterozygosity across their genomes [17]. Our laboratory has recently used the DO mouse population to conduct a behavioral and expression genetics analysis of a model for progressive ethanol consumption using prolonged intermittent voluntary 3-bottle choice drinking. These studies have identified multiple significant and suggestive behavioral quantitative trait loci (QTL) associated with various aspects of ethanol drinking behavior [18].

Here we present an in-depth analysis of transcriptomic responses in prefrontal cortex from the above DO study of intermittent ethanol consumption. Using multiple network analysis methods, along with expression QTL mapping and differential expression analysis, we have identified several groups of co-expressed genes that correlated strongly with ethanol consumption, and were overrepresented in genes differentially expressed between high and low ethanol consuming mice or control animals. Genes within these groups, particularly those identified as drivers of gene expression based on connectivity, had significant cis eQTLs, indicating genetic influence on gene expression. Biologically, one of the most notable findings was the abundance of myelin structural and myelin regulatory genes, indicating that regulation of myelination may be a highly conserved PFC response to prolonged ethanol exposure, and that this response is influenced by genetic variation. Regulators of such network responses, which we have identified as candidate genes, may underlie the observed behavioral phenotypes such as escalation of ethanol consumption [18].

## Materials & Methods

### Ethics Statement

All animal care and euthanasia procedures were performed in accordance with the rules and regulations established by the United States Department of Agriculture Animal Welfare Act and Regulations, Public Health Services Policy on Humane Care and Use of Laboratory Animals, and American Association for Accreditation of Laboratory Animal Care.

### Animals

Male Diversity Outbred (DO) mice (n=636) were acquired from Jackson Laboratories at 4 weeks of age, in 7 cohorts of 106 mice on average, spanning DO generations 21-24. Mice were singly housed in temperature and humidity-controlled vivariums on cedar shaving bedding with alternating 12 hour light/dark phases. At all points during the study mice were given free access to water and standard rodent chow (#7912, Harlan Teklad).

### Intermittent Ethanol Access

Mice received ethanol through intermittent access 3-bottle choice drinking. On Mondays, Wednesdays, and Fridays, for a period of 4 weeks, mice were given bottles with 15% v/v ethanol, 30% v/v ethanol, or water for a 24-hour period. Bottle order on the cages was randomized in order to avoid any effects from positional preference. On each drinking day, bottles were placed at the beginning of a dark phase (Figure 1a), and the amount consumed was measured at the end of the following light phase. Control mice were given 3 bottles of water. To account for evaporation, bottles were placed in an uninhabited control cage and the amount of fluid loss was subtracted from each day’s readings. Total ethanol consumption was calculated as amount 15% ethanol consumed (g/kg) + amount 30% ethanol consumed (g/kg). Total ethanol preference was represented by the fraction, in mL, of total ethanol (mL(15%EtOH) + mL(30%EtOH)) over mL total fluid (mL(water + mL(15%EtOH) + mL(30%EtOH)) intake.

**Figure 1:**
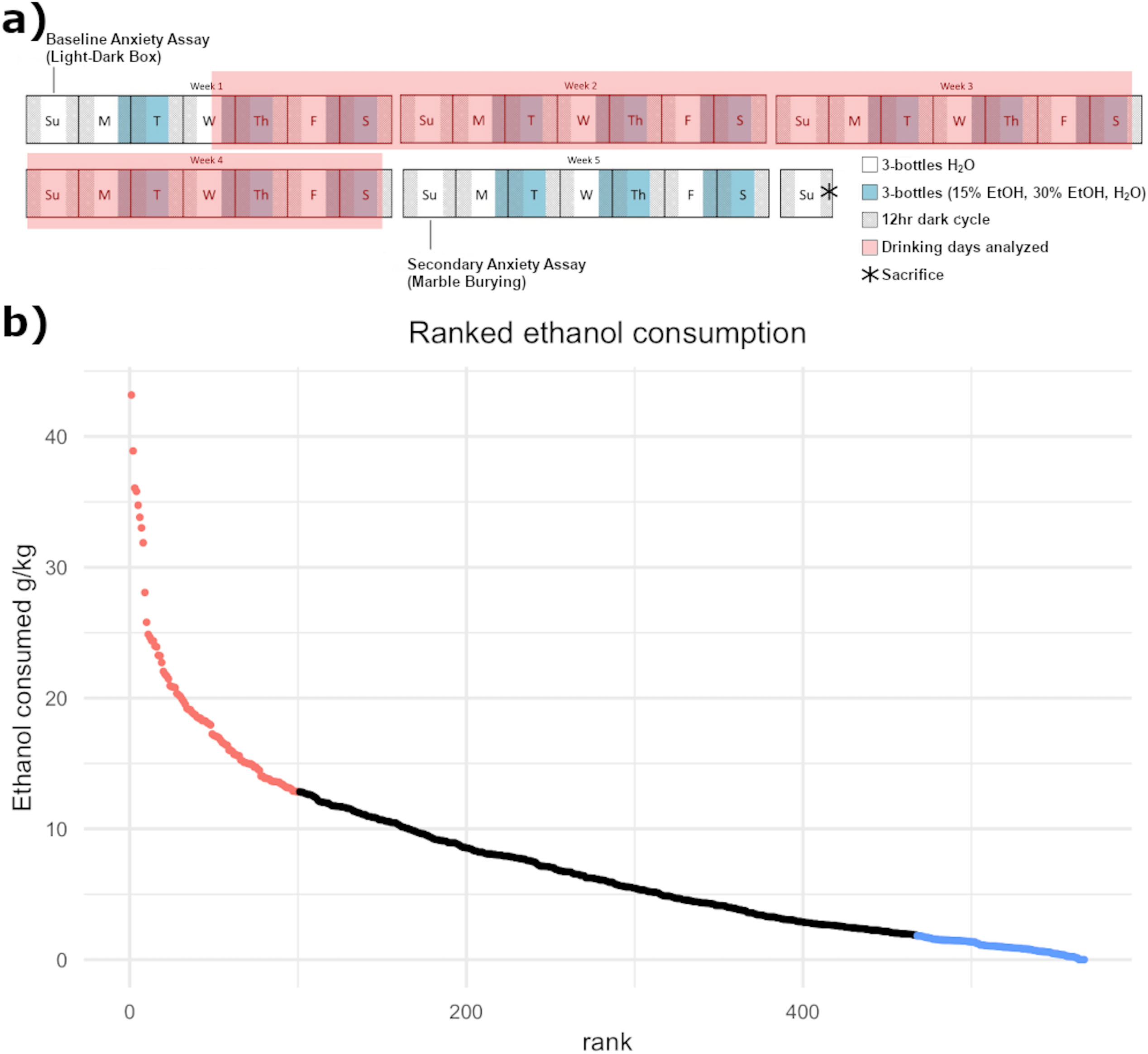
**a)** Overview of Intermittent Access Ethanol study with drinking days highlighted. **b)** Daily ethanol consumption for each mouse, in g/kg, ranked highest to lowest with 100 highest and lowest drinking mice that were selected for RNAseq highlighted in red and blue.

### Tissue processing, Genotyping, and RNAseq

Mice were sacrificed by cervical dislocation 24 hours after their final drinking session. Nine brain-regions (prefrontal cortex, nucleus accumbens, caudate, septum, amygdala, cerebellum, entorhinal cortex, hypothalamus, and ventral midbrain) were immediately dissected by punch microdissection based on stereotactic coordinates provided by the Allen Brain Atlas [19]. Tail tip samples were also taken for genotyping. Tissue was immediately flash frozen in liquid nitrogen and stored at -80° C.

Genotyping was performed by NeoGen Inc (Lincoln, NE) using the GigaMUGA microarray (N SNPs = 141,090; N CNVs = 2,169), which has markers optimized to be informative across the 8 DO founder strains. Mice were genotyped in two batches (N Batch 1 = 210, N Batch 2 = 420). Prefrontal cortex samples from 220 mice (100 highest drinking, 100 lowest drinking, and 20 control) were chosen for RNA-seq analysis based on average total ethanol consumption values during the fourth week of ethanol access. RNA was extracted and purified using RNeasy mRNA kits (Qiagen, Germantown, MD). Quality of isolated RNA was assessed using a NanoDrop (Thermo Fisher Scientific, Waltham, MA) and BioAnalyzer 2100 (Agilent Technologies, Santa Clara, CA). All samples had RIN values exceeding 8, with the exception of only 2 samples that had RIN values between 7 and 8. Samples were shipped on dry ice for poly-A selected library construction and sequencing at the Novogene Bioinformatics Institute (Sacramento, CA) as paired-end sequence fragments (150 bp) on the Illumina HiSeqX system resulting in approximately 20 million reads/sample.

### RNAseq Data Processing

Raw read counts were assessed for quality using FastQC (v0.11.7) by base sequence quality, tile sequence quality, base sequence content, GC content, sequence length, sequence duplication levels, and adapter content. Trimmomatic (v0.38) was then used to trim adapters and poor-quality bases at the 5’ and 3’ end of each read. Alignment and count generation was done by genome reconstruction by RNAseq (GBRS), a method that has been optimized to align RNAseq data to DO founder strains [17]. Transcripts with consistently low read counts across the DO mouse individuals were removed from further analysis (median total count < 1). Data were further examined for batch effects by principal component analysis. Principal component analysis was performed on transcripts per million (TPM) values.

### Differential Expression Analysis and Correlation to Ethanol Consumption

Differential expression (DE) analysis was run using the DESeq2 package for R [20]. In accordance with DESeq2 requirements, analysis began with raw read counts obtained from GBRS. The model was based primarily on drinking group, and included cohort as a covariate based on PCA analysis. Differential gene expression between high drinking and control mice was the comparison of interest for this report. P-values were adjusted for multiple testing using the Benjamini and Hochberg method. Among the DE results obtained, some FDR adjusted p-values appeared as NAs in the final results (Suppl. Table 1). The DEseq2 manual states that this may occur for genes which have zero or extremely low counts, or genes which extreme outliers [20]. RNAseq results were filtered for low counts, though the majority of genes that appeared as NAs had a lower base mean count (<15), indicating that these genes passed a low count filter of median total count < 1 but were still low expression compared to other genes in the dataset. FDR adjusted p-values ≤ 0.1 were considered significantly differentially expressed in further analyses.

Individual gene expression variance stablilized TPM counts were also correlated to ethanol consumption and preference using Spearman rank correlation with correction for multiple testing using the Benjamini and Hochberg method. Correlation corrected p-values ≤ 0.05 were considered significant.

### Network Analysis

Network analysis on RNAseq data was done using Weight Gene Correlated Network Analysis (WGCNA) [21]. ComBat for RNAseq, known as ComBat-*seq*, was used to account for the batch effect of breeding cohort [22]. TPMs were then calculated from raw values after ComBatSeq. Biweight mid-correlation and topological overlap mapping was used to calculate network adjacency. The treeCut() command was then used with deepSplit values 0-4. DeepSplit = 3 was chosen based on multi-dimensional scale plots (Suppl. Figure S1). Module-membership scores and gene-trait correlations (ethanol consumption and preference) were run for each gene. Module eigengenes (MEs), representing variation within each module, were also correlated to ethanol consumption and preference using Spearman rank correlation. Module connectivity metrics, including total network connectivity (kTot) and intramodular connectivity (kIn) were calculated in order to prioritize hub-genes within modules.

In order to identify key drivers within WGCNA modules, Bayes network learning using max-min Hill Climbing and Key Driver Analysis (KDA) [23, 24] was performed using the bnlearn package for R (https://www.bnlearn.com/). KDA identifies two types of nodes of within a network: key drivers and hubs. Key drivers are defined as nodes with a greater number of neighbors (nodes that have an edge signifying an interaction) than the mean number of neighboring nodes across the network plus the standard deviation of neighboring nodes across the network, and hubs are nodes with a greater number of out-degrees (interactions where the node acts on a neighboring node rather than being acted upon) than the mean number of out-degrees plus 2X the standard deviation of out-degrees [24]. This approach has been used in a previous study of bone density in DO mice [23] that we adapted for our analysis.

### Bioinformatics

Biological pathway enrichment, using the ToppFun tool available from the ToppGene Suite (https://toppgene.cchmc.org/), was run on each WGCNA module, as well as the list of all DE genes, and on genes within trans bands identified by eQTL mapping. Functional enrichment for GeneOntology categories, human phenotypes, mouse phenotypes, drugs, disease states, and regulatory miRNAs was included. Minimum and maximum overlap for enrichment was set at 3 ≤ x ≤ 5000 genes. Multiple testing correction was done using the Benjamini and Hochberg method. Categories with overlap FDRs ≤ 0.05 were considered significant. Gene Ontology (GO) categories were then reduced for complexity using the semantic similarity analysis tool REVIGO (http://revigo.irb.hr/) [25]. The Gene Interactions database tool, available through the UCSC Genome Browser (http://genome.ucsc.edu), was used to identify interacting proteins with the products of candidate genes identified by network analysis and eQTL mapping [26, 27]. GWAS Catalog (https://www.ebi.ac.uk/gwas/home) was used to determine whether candidate genes had previously been associated with alcohol or any other drugs of abuse in previous human GWAS [28].

Potential effects of cell-type were accounted for using Fisher’s Exact test with built-in WGCNA functions. The Zeisel *et al*. datasets for gene expression based on neurological cell-type were used as reference datasets [29].

The GeneInteractions tool, available through the UCSC Genome Browser (http://genome.ucsc.edu), was used to identify interacting proteins with the products of candidate genes identified by network analysis and eQTL mapping.

### Expression QTL (eQTL) Mapping

Read counts generated by GBRS were normalized with variance stabilizing transformation (VST) using the DESeq2 package. Analysis of eQTL was then done using the qtl2 package [30]. VST normalized expression data was used as the phenotype input. The genome scan was performed using linear mixed models with a kinship matrix to account for relatedness among individuals. Haplotype probabilities and breeding cohorts were also included as covariates. P-values were generated from LOD scores. In order to determine LOD thresholds for significant and suggestive QTLs, permutation analysis was run on ∼1% of genes randomly selected from genes from the RNAseq data (n=220). Permutation analysis and LOD thresholds for p-values of 0.63 (suggestive) and 0.05 (significant) were run using functions within in qtl2 package. Confidence intervals for eQTLs were defined as the region ± 1.5 LOD from the peak marker. Cis-eQTLs, loci for which genetic variation is associated with expression changes in local genes, were of particular interest to our analyses. To identify cis-eQTLs in the dataset: 1) the eQTLs physical position needed to be within 2Mb of the associated gene’s physical position and 2) the confidence interval of the eQTL needed to be less than 2 Mb in size. Trans bands were regions of the genome where several distant genes had eQTLs. Trans band boundaries were defined by obtaining confidence intervals (CIs) for all eQTLs in the region using 1.5 LOD drop. Regions of the genome that were included in CIs for more than 10% of the trans eQTLs linked to the region were considered within the trans band.

The Ensembl online genome database was used for gene annotation. 55,418 known mouse genes were retrieved from this database. Due to occasional updating of the Ensembl database that results in genes being removed, split, or renamed entirely, there were 384 genes in the GBRS eQTL dataset that were missing from the Ensembl annotations. Any eQTLs correlated with these missing genes were removed from further analysis. Best Linear Unbiased Predictors (BLUPs) were then used to estimate QTL haplotype effects at each locus of interest as this method is useful for determining QTL effects in multi-parent populations [30].

Eigengene expression QTL mapping was also run using the eigenvalues (known as eigengene by WGCNA convention [21]) to represent composite group expression of modules identified by WGCNA. EQTL analysis of eigengenes was run with the qtl2 package using linear mixed models with a kinship matrix, and haplotype probabilities and breeding cohorts were also included as covariates as described above.

### Candidate Gene Prioritization

To prioritize candidate genes that were most likely to influence increased ethanol consumption across the DO mouse panel, each gene’s correlation to ethanol consumption, whether they had a significant expression QTL, differential expression between High Drinkers and Controls, presence in a WGCNA module with significant correlation to drinking, whether or not the gene was a key driver in the module’s Bayes inferred network, presence of the gene within a Gene Ontology category for a process known to be involved in the brain’s ethanol response, and if that gene had previously been identified in a GWAS study of alcohol use disorder were all taken into account. Two genes that were filtered for low expression before WGCNA analysis (*Ccdc129* and *Shroom3*), but were included in eQTL mapping, were eliminated from candidate gene prioritization.

### Data Sharing

Transcriptomic data obtained from RNAseq are publicly available through the Gene Expression Omnibus (https://www.ncbi.nlm.nih.gov/geo/).

## Results

### Intermittent Access Ethanol

636 DO mice were used for intermittent access ethanol drinking by 3-bottle choice for 11 drinking days over 4 weeks (Figure 1a). As discussed in a previous publication, these mice significantly increased the total amount of ethanol they consumed over the course of the study [18]. Of these, 200 mice were selected for RNAseq analysis, as RNAseq on 636 mice would have been cost prohibitive. Mice were selected based on mean daily consumption (in g/kg) across all drinking days. The 100 highest (High Drinkers) and 100 lowest (Low Drinkers) drinking mice were selected for RNAseq because these were most likely to show differences in gene expression and in genetic variation underlying gene expression differences (Figure 1b). Additionally, 20 of the control 49 mice, that drank only water, were randomly selected for inclusion in RNAseq analysis. This brought the total number of RNAseq samples run to 220.

### RNAseq Quality Assessment

FastQC analysis showed high overall quality scores across samples. Most samples were comparable in GC content, per base N content, sequence duplication levels, and adapter content. Only a few samples indicated low sequence counts. These samples were sent for resequencing and results were merged with original sequencing results. The final counts and quality scores of merged sequencing runs were comparable to other samples. As is typical of RNAseq analysis, base quality values at the far ends of each read were lower than the rest of the fragment. After 5’/3’ trimming with Trimmomatic, all samples passed quality control and were included in downstream analyses.

Transcripts per million (TPM) were generated for differential gene expression (DE) and network analyses. PCA analysis of TPM values showed a very mild batch effect of breeding cohort. For this reason, cohort was included as a covariate in DE analysis with DESeq2, and, for network analysis, was corrected using the ComBat-*seq* method [22].

### Differential Gene Expression

A total of 47,643 transcripts were aligned with GBRS. For differential gene expression analysis, TPM values were generated and then log-scaled. DEseq was used to compare gene expression between High Drinkers, Low Drinkers, and Control mice. At a false discovery rate of 0.10, 8705 genes were differentially expressed between High Drinkers and Low Drinkers, 677 between High Drinkers and Controls, and 1294 between Low Drinkers and Controls (Figure 2a-c, Suppl. Table S1).

**Figure 2:**
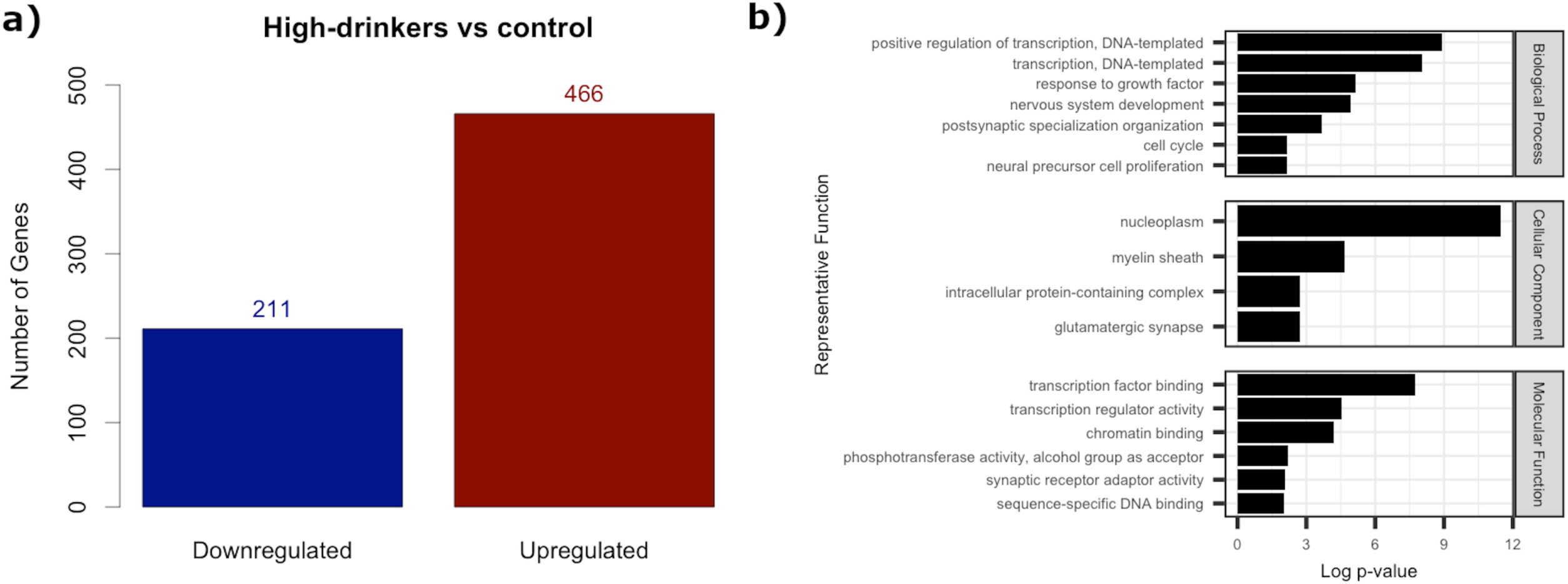
**a)** Number of genes upregulated and downregulated at FDR ≤ 0.1 between High-Drinkers and Low-Drinkers. **b)** Main Gene Ontology categories, paired by semantic similarity, represented by genes significantly differentially expressed at FDR ≤ 0.1 between High-Drinkers and Control.

The greatest number of DE genes were seen between Low Drinkers and High Drinkers. These genes were enriched for Gene Ontology categories related to cell-cycle regulation, protein localization and transport, organization of cell projections, catalytic complex components, nucleoplasm and organelle envelope components, regional organization of the plasma membrane, chromatin binding, myelin sheath components, nervous system development, and regulation of neuronal cell death (Suppl. Figure S2-S3, Suppl. Table S3). Between Low Drinkers and Control mice, differentially expressed genes represented Gene Ontology categories involved in energy coupled proton transport, cellular respiration, mitochondrial components, ribosomal components, as well as myelin sheath and synaptic vesicle components (Suppl. Figure S2-S3, Suppl. Table S3). Between High Drinkers and Controls, 466 were upregulated and 211 were downregulated (Figure 2a). Bioinformatically, these genes were enriched for Gene Ontology categories involved in regulation of DNA transcription, nervous system development, myelination, cell cycle regulation, synaptic activity, chromatin binding, and growth factor response (Figure 2d, Suppl. Table S3). Genes differentially expressed between High Drinkers and Controls were of most interest because these genes are likely to be those whose expression is either modulated by prolonged ethanol drinking or contributed to an increased drinking phenotype.

### Expression QTL Mapping

Expression QTL mapping resulted in a total of 16,964 significant eQTLs following filtering for LOD scores > 8.45 (equivalent to p-value ≤ 0.05). Of these, 9,515 were cis-eQTLs representing 8,090 genes (Figure 5a-b, Suppl. Table S4). Chr 11 had the largest number of total eQTLs, but Chr 7 had the largest number of cis eQTLs and Chr 18 had the highest number of trans eQTLs (Figure 5a-b, Suppl. Figure S6, Suppl. Table S3).

Additionally, trans-bands were identified on Chr 2 and Chr 13 and Chr 18 (Figure 5b, Suppl. Figure S7). These trans-band loci were associated with expression regulation of multiple remotely located genes having possible related functions or mechanisms of regulation. Biologically, both trans bands on Chr 18 were enriched for synapse organization, neurotransmitter transport and release, and nervous system development. These regions of Chr 18 were also significantly enriched for genes associated with cocaine, alcohol, methamphetamine abuse (Suppl. Table S6, Suppl. Figure S8-S9). The trans band on Chr 13 was also enriched for dopamine signaling at the synapse, along with locomotory behavior and amphetamine response (Suppl. Table S6, Suppl. Figure S10). This finding is, perhaps, not surprising as dopamine signaling has documented involvement in locomotor activity and ataxia, and in the rewarding effects of drugs of abuse [31–33]. Our findings not only reinforce such previous observations, but also suggest that Chr 13: 112.83-119.48 Mb and Chr 18: 71.24-73.73, Chr 18: 37.34-45.63 may code for important molecular regulators of synaptic signaling in response to ethanol.

### Co-Expression Networks Within the PFC

WGCNA was used for network analysis to identify co-regulated genes of possible related biological function. Log-scaled TPM values, corrected for cohort using ComBat-*seq*, were used for network analysis because this normalization method fit scale-free topology (one of the assumptions of WGCNA) better than Variance Stabilizing Transformation (Suppl. Figure S4a-b). A soft-thresholding power of 4 was selected for network construction. This process identified 31 co-expression modules ranging in size from 33 to 10,298 genes (Table 1). 20 modules showed an overall correlation of expression, as measured by Spearman rank, to ethanol consumption or ethanol preference. 14 of these were negatively correlated and 6 were positively correlated. All modules that significantly correlated to consumption were also significantly correlated, in the same direction, to ethanol preference (Figure 3). Of these, 7 modules were significantly enriched for genes that were differentially expressed between High Drinkers and Controls (Table 2).

**Figure 3:**
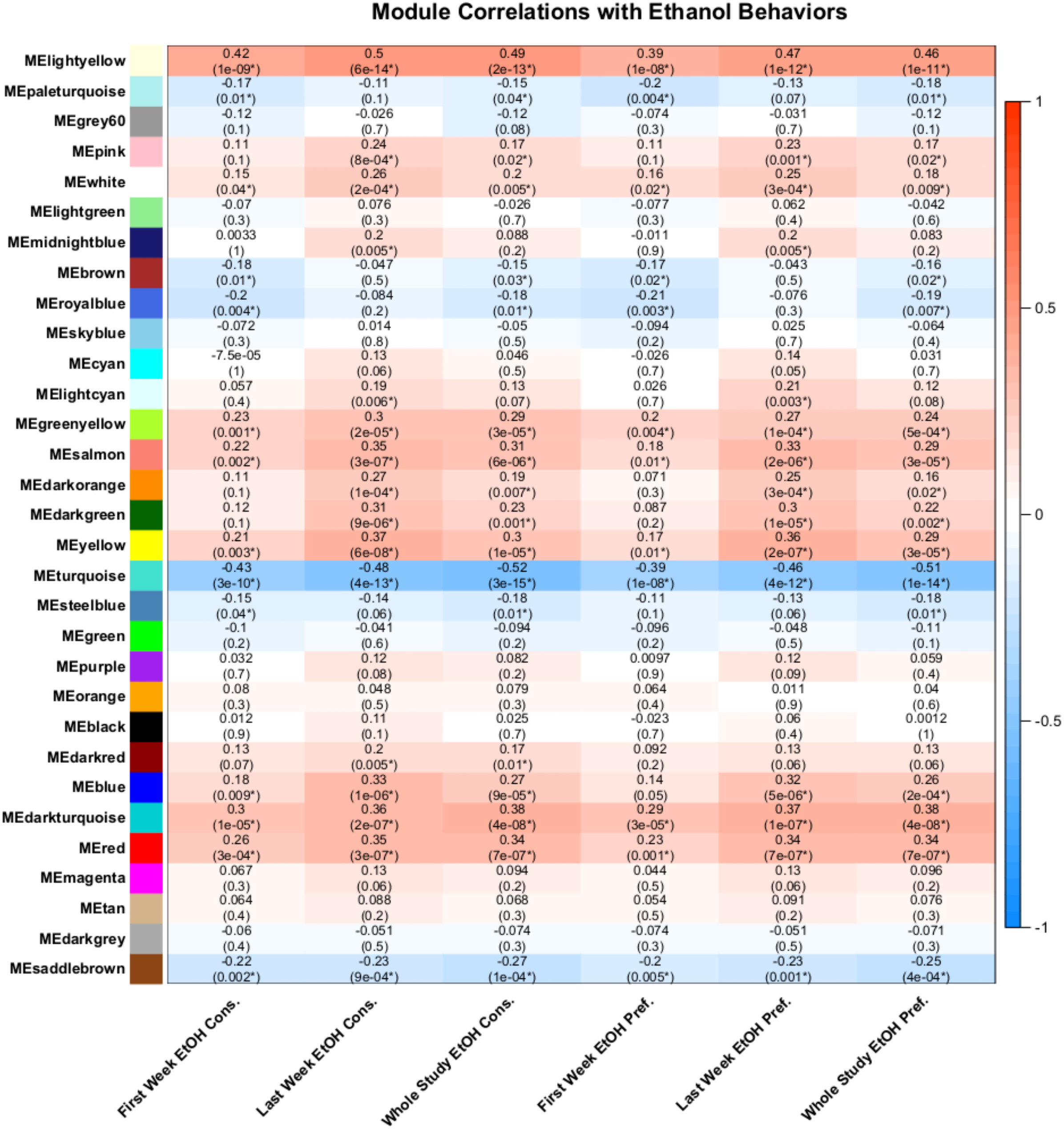
Heatmap of correlation between ethanol consumption and ethanol preference and the module eigengene of each WGCNA module.

**Table 1:** WGCNA module sizes.

| **Module** | **Number of Genes** | **Module** | **Number of Genes** |
| --- | --- | --- | --- |
| black | 375 | midnightblue | 89 |
| blue | 1,921 | orange | 53 |
| brown | 1,611 | paleturquoise | 33 |
| cyan | 91 | pink | 171 |
| darkgreen | 62 | purple | 140 |
| darkgrey | 54 | red | 558 |
| darkorange | 51 | royalblue | 70 |
| darkred | 69 | saddlebrown | 35 |
| darkturquoise | 55 | salmon | 98 |
| green | 611 | skyblue | 39 |
| greenyellow | 106 | steelblue | 34 |
| grey60 | 78 | tan | 104 |
| lightcyan | 84 | turquoise | 10,298 |
| lightgreen | 77 | white | 50 |
| lightyellow | 71 | yellow | 774 |
| magenta | 144 |  |  |

**Table 2:** Overlap between WGCNA modules and ethanol consumption regulated genes.

| Module | Number of DE genes/Module Size | P-value |
| --- | --- | --- |
| greenyellow | 45/106 | $2.41 \times 10^{-34}$ |
| red | 87/558 | $7.74 \times 10^{-29}$ |
| salmon | 38/98 | $4.58 \times 10^{-27}$ |
| blue | 126/1,921 | $9.35 \times 10^{-9}$ |
| yellow | 59/774 | $1.40 \times 10^{-5}$ |
| darkorange | 11/51 | $2.50 \times 10^{-4}$ |
| darkgreen | 10/62 | 0.0105 |

WGCNA modules having significant overlap with differentially expressed genes in the DESeq2 analysis of LWECD vs. control animals are shown. P-values reflect Fisher's exact test for over-representation.

**Table 3:** eQTL content across chromosomes.

| **Chromosome** | **Total significant eQTLs** | **Number of cis eQTLs** | **Number of trans eQTLs** |
| --- | --- | --- | --- |
| **1** | 1080 | 669 | 411 |
| **2** | 1412 | 715 | 697 |
| **3** | 946 | 543 | 403 |
| **4** | 1110 | 667 | 443 |
| **5** | 1188 | 714 | 474 |
| **6** | 854 | 529 | 325 |
| **7** | 1359 | 858 | 501 |
| **8** | 701 | 441 | 260 |
| **9** | 855 | 545 | 310 |
| **10** | 664 | 379 | 285 |
| **11** | 1232 | 736 | 496 |
| **12** | 632 | 319 | 313 |
| **13** | 594 | 307 | 287 |
| **14** | 608 | 282 | 326 |
| **15** | 534 | 322 | 212 |
| **16** | 408 | 242 | 166 |
| **17** | 806 | 492 | 368 |
| **18** | 1118 | 264 | 854 |
| **19** | 378 | 277 | 101 |
| **X** | 430 | 214 | 216 |

Table displays number and type of eQTLs per chromosome using a $FDR \leq 0.05$ for identification of significant eQTL peaks.

We generated module eigengenes (first principal component of genes within a module) of WGCNA modules to enable more facile module correlation with behavioral data as well as identification of genetic regulators of WGCNA modules. Eigengenes of each expression module identified by WGCNA were used as phenotypes in Rqtl2 analysis. This approach identified 3 eigengene QTLs (egQTLs) that were significant at a LOD score of ≥ 9.66 (equivalent to a p-value of 0.05), and 45 QTLs with a suggestive LOD score (equivalent to a p-value of 0.63) (Suppl. Table S7). Of these, 2 significant egQTLs were associated with the paleturquoise module, a module with relatively low correlation to total ethanol intake, though this correlation did reach significance to first week and whole study consumption and preference (Figure 3). These egQTLs were found in non-overlapping confidence intervals on Chr 4 (Suppl. Table S7). The third egQTL was present on Chr 13 and was associated with the pink module (Suppl. Figure S11, Suppl. Table S7), a module that had somewhat lower, though still significant, correlation to last week and whole study consumption and preference (Figure 3). Among modules with suggestive QTLs, the salmon, greenyellow, turquoise, and dark turquoise modules were among those with the highest correlation to ethanol intake. The red and lightyellow modules, in spite of high correlation with ethanol intake, were not associated with any suggestive or significant eigengene QTLs.

Since we used whole tissue samples, cell-type enrichment was performed to determine whether cell population over-representation was driving co-expression in certain WGCNA modules. Based on a published single-cell RNAseq from mouse somatosensory cortex and hippocampus as the reference dataset [29], only two modules showed enrichment for a specific cell-type. The orange module was significantly enriched for oligodendrocytes (Suppl. Figure S5). This was an unsurprising finding, as the orange module contained classic myelin component genes such as myelin basic protein (*Mbp*), myelin oligodendrocyte associated basic protein (*Mobp*), myelin associated glycoprotein (*Mag*), myelin and lymphocyte protein (*Mal*), and proteolipid protein 1 (*Plp1*). *Mbp* was also a key driver in the orange module as assessed by Bayes network analysis (Suppl. Table S5). None of these genes had significant eQTLs, however, some of their known interacting proteins, identified with the UCSC Gene Interactions tool, such as *Myd88*, *Fyn*, *Diaph1*, *Tirap*, and *Ticam1*, had significant trans eQTLs (Suppl. Table S4). The black module, however, was significantly enriched for mural, endothelial, and ependymal cells (Suppl. Figure S5). Endothelial cells represent the inner cell layer of blood vessels, and mural cells include vascular smooth muscle cells and pericytes, and ependymal cells are found in the epithelial lining of the ventricular system in the brain. These results suggest that genes responsible for maintaining the circulation of blood and cerebrospinal fluid in the brain are co-expressed in the black module. However, neither of these modules were significantly correlated to ethanol consumption, nor were they significantly enriched in genes that were differentially expressed between High Drinkers and Controls (Figure 3, Table 2). Drinking Correlated Modules Enriched for DE genes

#### Salmon Module

The salmon module was one of the modules that was most significantly correlated to last week ethanol consumption (r^2^ = 0.35, p-value = 3 x 10^-7^), whole study ethanol consumption (r^2^ = 0.31, p-value = 6 x 10^-6^), last week ethanol preference (r^2^ = 0.33, p-value = 2 x 10^-6^), and whole study ethanol preference (r^2^ = 0.29, p-value = 3 x 10^-5^) (Figure 3). Not only was the module eigengene of the salmon module significantly correlated to ethanol consumption, individual genes within this module also showed a positive correlation between module membership, which uses the correlation between each gene and the module eigengene to measure the fit of each gene to the module, and correlation to ethanol consumption (Figure 4a). The salmon module also contained a significant number of genes that were differentially expressed between High Drinker and Control mice (p-value = 4.58 x 10^-27^) (Table 2). GO analysis for this module showed enrichment for categories related to chromatin binding, regulation of transcription and transcription co-regulators, protein deacetylation, neurogenesis, neuron differentiation and development, regulation of post synaptic potential, axonogenesis, development of thorny excrescence, and dendritic spine development (Figure 4b, Suppl. Table S3). Within these categories, Bayes network analysis showed that *Acin1* was a significant hub gene, and that *Srf, Baiap2, Sin3b, Pbx2, Socs7, and Pygo2* were key drivers (Suppl. Table S5). Of these, only *Pygo2* had a significant eQTL, a trans eQTL on Chr 18 (Suppl. Table S4). There were, however, known inhibitors and co-factors of some of these key drivers that did have significant eQTLs. These include, for example, *Srfbp1*, *Baiap2l1, Il6st, Il12rb1, Stat6, Lifr, Ifnar1, Ifnar2,* and a number of *Fox* family genes (Suppl. Table S4). These genes were also in GO categories related to neurogenesis. Further, all of the other identified key drivers, with the exception of *Tmem28* and *Cecr6* were in GO categories involved in regulation of transcription and translation. Genes in these categories included *Dpf1, Mafg, Sin3b, Hdac5, Srf,* and *Pbx2*, all of which have documented roles in regulation of synaptic transmission in the brain [34, 35], particularly via regulation of chromatin binding [36], further indicating that this module represents a synaptic signaling module. *Srrm3* was another significant hub gene within this module and, while not in any significantly enriched GO categories, this gene was regulated by several miRNAs whose targets significantly overlapped with the salmon module (Suppl. Table S3, S5). These findings suggest that expression of genes within this module are modulated by heavy ethanol drinking, and that heavier drinking mice tend to show higher expression of genes within this module (Figure 4a). Further, GO analysis indicates that these genes are involved in the regulation of expression, at the gene and protein level, of genes involved in the development of presynaptic projections and downstream modulation of postsynaptic signaling (Suppl. Table S3).

**Figure 4:**
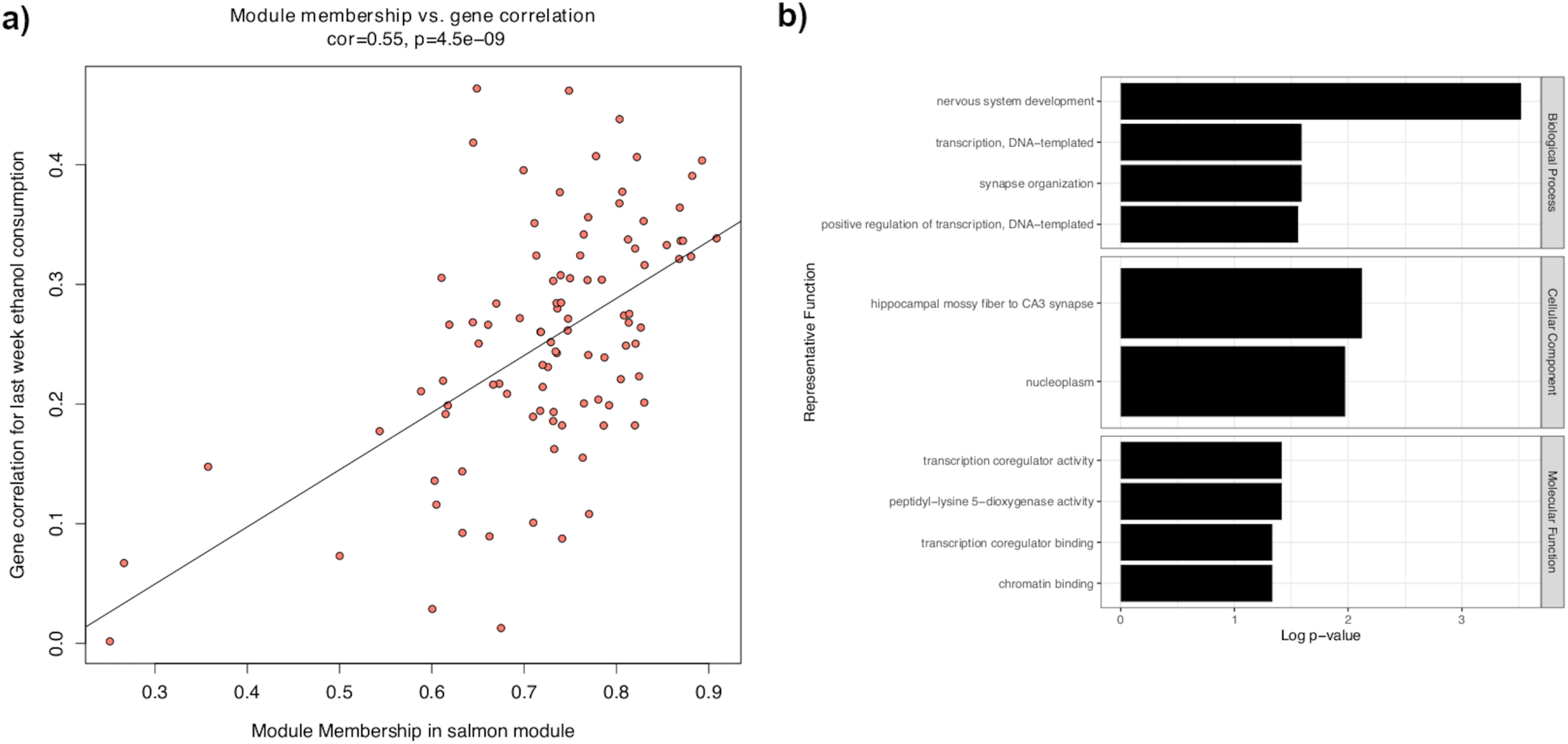
**a)** Scatterplot of module membership score (correlation between gene and the module eigengene) vs gene correlation with ethanol consumption across the whole study. **b)** Main Gene Ontology categories, paired by semantic similarity, represented by genes in the salmon module.

**Figure 5:**
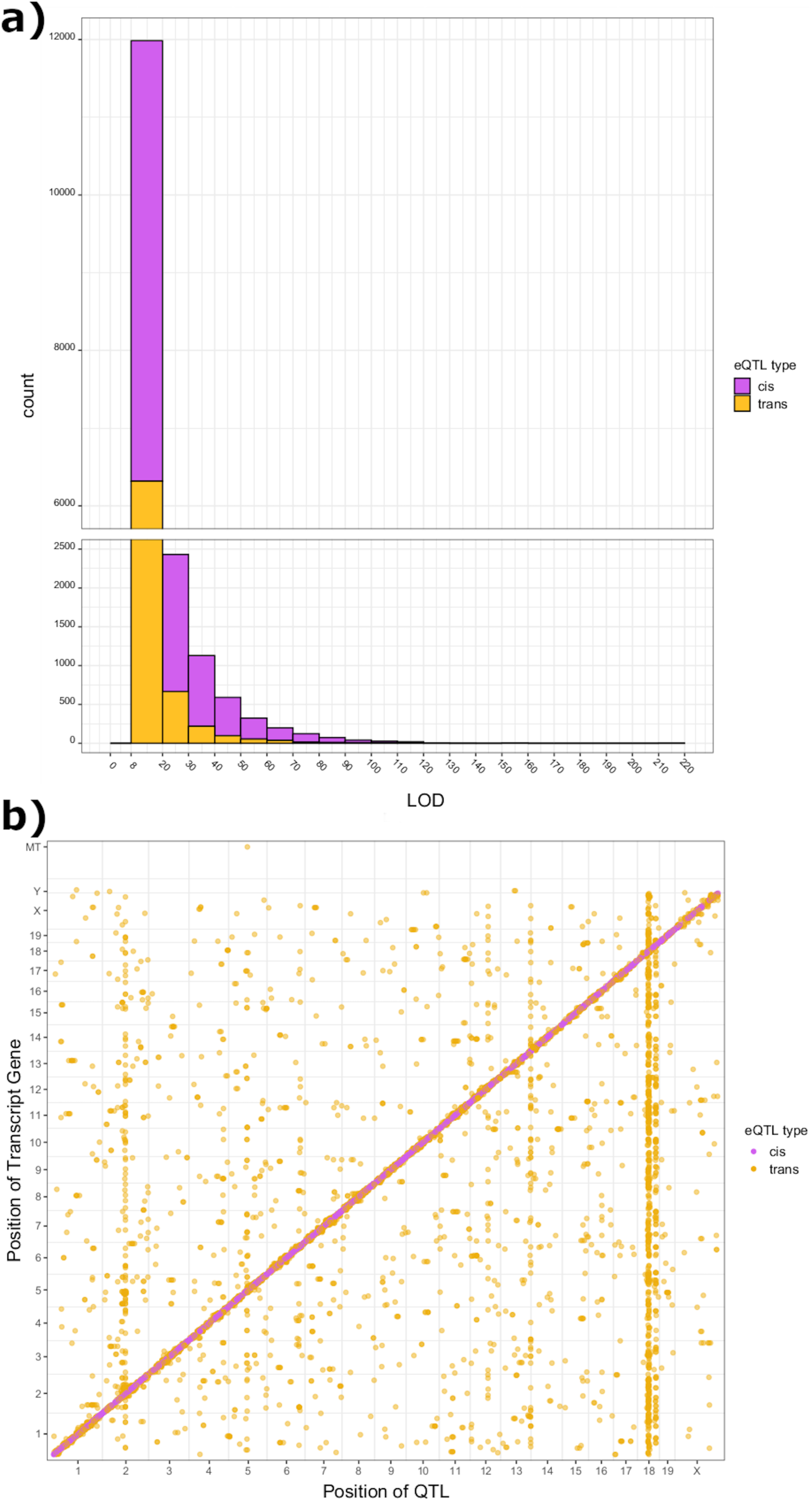
**a)** Bar plot of all expression QTLs with LOD scores > 8.45, a threshold determined to be equivalent to a p-value of 0.05. **b)** Location of all expression QTLs with LOD scores > 8.45.

#### Greenyellow Module

The greenyellow module was also significantly correlated to both ethanol consumption and ethanol preference (last week ethanol consumption: r^2^ = 0.3, p-value = 2 x 10^-5^, whole study ethanol consumption: r^2^ = 0.29, p-value = 3 x 10^-5^, last week ethanol preference: r^2^ = 0.27, p-value = 1 x 10^-4^, whole study ethanol preference: r^2^ = 0.24, p-value = 5 x 10^-4^) (Figure 3), and was significantly enriched for genes that were differentially expressed between High Drinker and Control mice (Table 2). Bioinformatically, this module was highly enriched for GO categories related to DNA binding, regulation of transcription factor binding, regulation of transcription factor activity (particularly RNA polymerase II), regulation of protein kinase activity, regulation of various metabolic processes, regulation of hormone response pathways, and regulation of apoptosis. Several of these categories included *Fos* and *Fos*-interacting genes, indicating this module was enriched in immediate-early genes (Suppl. Table S3). Interestingly, however, few of these genes were identified as key drivers or hubs in this module (Suppl. Table S5).

#### Turquoise Module

The largest module identified in this dataset was the turquoise module. In previous studies from our group that used WGCNA, the turquoise module was often large [14, 15, 37]. The turquoise module identified from DO mice was very large at over 10,000 genes (Table 1). However, this module was both significantly enriched for DE genes between High Drinker and Control mice (Table 1), and highly negatively correlated with ethanol consumption and ethanol preference (first week ethanol consumption: r^2^ = -0.43, p-value = 3 x 10^-10^, last week ethanol consumption: r^2^ = -0.48, p-value = 4 x 10^-13^, whole study ethanol consumption: r^2^ = -0.52, p-value = 3 x 10^-15^, first week ethanol preference: r^2^ = -0.39, p-value = 1 x 10^-8^, last week ethanol preference: r^2^ = -0.46, p-value = 4 x 10^-12^, whole study ethanol preference: r^2^ = -0.51, p-value = 1 x 10^-14^) (Figure 3). Unsurprisingly, this module was enriched for a wide variety of categories through GO analysis (Suppl. Table S3). Overall, these processes included cellular catabolic processes, cellular organization including organelle organization, cellular component biogenesis, cellular stress response, cell cycle regulation and cell growth, RNA and DNA binding, ubiquitin protein ligase binding, pyrophosphatase activity, transferase activity, GDP binding, nervous system development, nucleoplasm elements, nuclear envelope elements, intracellular protein-containing complexes, and components of cell junctions, cell projection, the plasma membrane, and the myelin sheath (Suppl. Table S3).

### Drinking Correlated Modules Not Enriched for DE genes

#### Lightyellow Module

Although many of the most highly drinking correlated modules were also enriched for DE genes between High Drinker and Control mice, there were also a few drinking correlated modules that were not enriched in DE genes. The lightyellow module was one of these. This module was very strongly positively correlated to ethanol consumption and ethanol preference (first week ethanol consumption: r^2^ = 0.42, p-value = 1 x 10^-9^, last week ethanol consumption: r^2^ = 0.50, p-value = 6 x 10^-14^, whole study ethanol consumption: r^2^ = 0.49, p-value = 2 x 10^-13^, first week ethanol preference: r^2^ = 0.39, p-value = 1 x 10^-8^, last week ethanol preference: r^2^ = 0.47, p-value = 1 x 10^-12^, whole study ethanol preference: r^2^ = 0.46, p-value = 1 x 10^-11^) (Figure 3). In terms of GO analysis, this module was significantly enriched for nucleotide biosynthetic processes, structural components of ribosomes such as *Rps* and *Rpl* genes, ion transport and ion binding, protein targeting and protein modification, and chaperone and other unfold protein binding. *Rpl6* and *Rpl36al* were also key drivers within this module (Suppl. Table S5). This module was also very enriched for genes involved in cellular respiration including NADH dehydrogenase, cytochrome-c, and other oxidoreductases. Lightyellow module genes that were found in these GO categories included *Cox6* and *Cox7* genes (Suppl. Table S3) and *Nduf* family genes, of which 5 were key drivers within the module (Suppl. Table S5).

#### Darkgreen Module

The darkgreen module was one of the smaller modules that was also significantly correlated to ethanol drinking (Figure 3) but did contain many genes that were differentially expressed between High Drinker and Control mice (Table 1). This module was strongly positively correlated to last week ethanol consumption and ethanol preference (last week ethanol consumption: r^2^ = 0.31, p-value = 9 x 10^-6^, last week ethanol preference: r^2^ = 0.30, p-value = 1 x 10^-5^) (Figure 3). Bioinformatically, this module was enriched for many GO categories involved in nucleotide binding, neuron development and differentiation, structural components of the synapse, and regulation of synaptic signaling. Genes within this module that overlapped with these GO categories included genes that specifically function in synaptic signaling by cytoskeleton-associated vesicle trafficking such as *Nav2*, *Bsn*, *Kif21b*, *Mpp2*, *Smarca4*, and *Myo18a* (Suppl. Table S3). *Smarca4* was also a key driver in the darkgreen module (Suppl. Table S5).

### Candidate Gene Prioritization

Using the parameters outlined in the Methods section, 177 genes were selected as potential candidate genes. The number of candidate genes per module tended to, generally, be proportional to the size of the module. The blue, turquoise and yellow modules had the lowest number of candidate genes. The brown module, which was the 3^rd^ largest WGCNA module, was an exception having only 8 candidate genes compared to 21 in the slightly larger blue module and 12 in the yellow module which had 837 fewer genes than the brown module (Suppl. Table S8, sheet 1). Gene Ontology analysis of all candidate genes together, regardless of module assignment, showed enrichment (FDR ≤ 0.05) for 5 GO categories: cellular components, intracellular protein complexes, catalytic complexes, myelin sheath, and cell projections (Suppl. Table S8, sheet 2).

## Discussion

Due to the strong heritable component seen with AUD, identifying the genetic underpinnings of the tendency toward escalating consumption is important for understanding the etiology of this disorder and the physiological changes that result from prolonged excessive ethanol intake. We have made use of the DO mouse panel with IEA as an animal model of prolonged voluntary ethanol consumption in a genetically diverse population to more accurately reflect mechanisms underlying AUD in the human population. We have recently shown that the IEA drinking paradigm leads to a progressive increase in consumption in DO mice over 4 weeks of drinking, and that variation in specific regions of the genome show significant association to the amount a mouse will consume [18]. Building upon that behavioral genetic analysis, here we have used network analysis methods and expression QTL mapping to characterize the gene expression changes that occur in the prefrontal cortex, the region of the brain responsible for impulse control and other executive functions, after 4 weeks of IEA. Through these analyses, we identified several sub-networks, known as modules, of co-expressed genes that were correlated with ethanol intake, and showed significant differential expression between High Drinker and Control mice. We have also identified several novel associations between genomic regions and gene expression through eQTL mapping, identified several potential candidate genes based on connectivity within co-expression modules that correlated to drinking and association with important genes involved in biological processes whose gene expression is highly correlated to drinking under IEA.

One notable observation from network analysis with WGCNA was that most modules correlated with ethanol intake were also significantly enriched for ethanol responsive genes. This finding is contrast to our previous transcriptomic studies after chronic ethanol exposure in monkeys and inbred mice which showed that co-expression modules in PFC tended to be either correlated to ethanol consumption or enriched in ethanol responsive genes, but not both [37]. That we have found modules of co-expressed genes that tend to both correlate to ethanol consumption and show significant differential expression, particularly between high drinking and ethanol naive mice, indicates that using a large and genetically diverse mouse model has perhaps allowed identification of transcriptomic responses in PFC that underlie environmental x genetic interactions in development of excessive ethanol consumption.

From these analyses, genes involved in regulation of myelination in the prefrontal cortex, and their interacting proteins, have emerged as significantly responsive to heavy ethanol consumption and as having significant eQTLs, suggesting genetic variance in their expression. Our group has previously reported decreased expression of myelin structural genes and myelin-interacting genes in the prefrontal cortex of mice following acute [31, 38] and chronic ethanol exposure in mice and macaques [15, 37, 39]. Other studies in animal models and human patients have also noted disruption of components of the myelin sheath at both the gene and protein level [40–43]. Reduced myelination in the frontal cortex has also been observed at the structural level in rodent models of fetal [44, 45], adolescent [46–48], and adult ethanol exposure [49, 50]. This disruption of cerebral myelination, and the expression of myelin regulatory genes following ethanol exposure, has also been associated with impaired memory and cognition [51] and lasting changes in ethanol sensitivity [38, 51]. From our analyses, certain myelin structural genes such as *Mbp, Mobp,* and *Mag* were significantly positively correlated with ethanol consumption. Of these, *Mbp* was a key driver of the orange module, and one of its interacting proteins (*Ticam1*) showed significant correlation to ethanol consumption or preference [18]. This an interesting finding, as previous studies have shown a reduction in myelin gene expression with ethanol exposure. Canonical myelin genes such as *Mbp* and *Mobp* showed no significant differences in expression between High Drinkers or Low Drinkers compared to Controls. However, there was a significant difference in *Mbp, Mag, Myd88, Fyn,* and *Ticam1* expression between High Drinkers and Low Drinkers, indicating a high degree of variance in the expression of these genes across DO mice (Suppl. Table S1). Expression QTL analysis, at both the gene and module level, however, indicated significant association between genomic variation and expression of myelin-regulatory genes (Suppl. Table S3, S7) and that these genes are key drivers of the co-expression of myelin genes (Suppl. Table S5). Our analyses of DO mice demonstrate that this modulation of myelination is a response that is observable across a genetically diverse population, and provides further evidence that ethanol-associated disruption of myelination may underlie the tendency toward heavy drinking and escalation of drinking that we have observed with prolonged exposure [18]. Our findings are particularly interesting because many of these myelin interacting genes are also interacting proteins for Toll-like receptor proteins or are adapter proteins in the Toll-like receptor/WNT signaling pathway [52, 53]. Several recent studies indicate that Toll-like receptors, particularly Toll-like receptor 4 (*Tlr4*), mediates ethanol’s effect on myelination in the prefrontal cortex [47, 52] and that modulation of *Tlr4* shows promise in attenuating reduced myelination and associated cognitive, anxiety-like, and social interaction symptoms in mice and rats exposed to ethanol *in utero* or during adolescence [53–56]. Our analyses were performed in adult DO mice, providing novel evidence that this approach may also be effective in reducing alcohol-associated demyelination, and resulting cognitive symptoms, in adults.

In addition to association between genotype and myelin regulatory genes, eQTL analysis at the gene and module level also identified several more novel candidate genes (Suppl. Table S3, S6-S7). The salmon module eigengene was strongly correlated with ethanol consumption and eigengene QTL analysis identified a suggestive egQTL on Chr 5 associated with the salmon module (Suppl. Table S7). Within this module *Pygo2* was a key driver gene (Suppl. Table S5) and had a significant trans eQTL on Chr 18 (Suppl. Table S4). This gene has several functions including chromatin binding, histone acetyltransferase and methyltransferase regulation, and WNT pathway regulation of beta-catenin signaling [57, 58]. This latter finding potentially has mechanistic implications on ethanol consumption, as the WNT signaling pathway is also regulated by *Gsk3β*, a gene that our group has previously shown to modulate ethanol consumption and withdrawal-induced anxiety in rodent models and to be associated with alcohol dependence in human genetic studies [59]. Interestingly, *Gsk3β* also had a significant trans eQTL on Chr 18 (Suppl. Table S7) having a support interval within 3.7 Mb of that for the *Pygo2* trans eQTL. This result indicates that this region of Chr 18 could influence ethanol phenotypes both via *Gsk3β* and the greater myelin response to ethanol.

Synaptic signaling and neurogenesis was a theme found in another WGCNA module (darkgreen) that correlated with drinking, was significantly enriched in DE genes (Table 2) and had a suggestive eigengene QTL on Chr 5 (Figure 3, Suppl. Figure S11). This module was enriched in GO categories involved in vesicle trafficking and neurodevelopment. In this module, *Arhgef18* was the only hub gene (Suppl. Table S5). This gene is a Rho guanine exchange factor involved in vesicle trafficking. This gene has also been shown to have higher expression in high-drinking rats [60]. In our analysis, this gene was also higher expression in High Drinker compared to Low Drinker DO mice (Suppl. Table S1). This gene, therefore, may represent another potential candidate gene that could be targeted to affect the gene expression response to prolonged ethanol drinking.

It is important to note that only male mice were used for these analyses due to the number of mice needed to achieve significant power to account for genetic effects and sex differences between males and females. The use of only male mice limits our ability to detect any sex-specific effect seen in the PFC gene expression response to prolonged intermittent ethanol drinking. However, future studies of the same design are targeted to study female mice. Based on previous studies of DO progenitor strains which found that female mice drank more than males with IEA [61], we expect that the magnitude of gene expression response to IEA may be greater in females, but that the same biological processes will be disrupted by several weeks of IEA. Further, based on previous studies of sex differences in ethanol intake in mouse models [62], female DO mice may also escalate their drinking to an even greater degree than we have observed in male mice.

Through use of genetic, transcriptomic, and bioinformatic analyses of Diversity Outbred mice, we have identified novel gene expression networks and potential regulatory mechanisms in prefrontal cortex that relate to progressive ethanol consumption. This approach identified disruption of PFC myelination as a conserved, highly significant response to ethanol that is influenced by genetic background. Our analyses also implicate ethanol’s effect on inflammatory response in the CNS, particularly via interleukin and Toll-like receptor signaling, as a regulator of the cerebral myelin response to prolonged ethanol. Other highly ethanol correlated and responsive processes included neurodevelopment, vesicle signaling at the synapse, and transcription regulation by chromatin modification. Further, expression QTL analysis identified a trans band on Chr 18 that was significantly associated with expression variation in several genes involved in *Wnt-Gsk3β* signaling, a pathway that we have previously implicated in PFC ethanol consumption. This led to several potential candidate genes as regulators of synaptic singling in response to ethanol including *Arhgef18, Smarca4, Baiap2l, Stat6,* and *Pygo2.* These pathways may also underlie escalation of ethanol consumption with prolonged high dose drinking seen in human AUD patients, making further exploration of their potential regulators important in the development of new therapeutics for dependent or problematic alcohol drinking.

## Acknowledgments

This work was supported by NIAAA grants P50AA022537 (MFM) and F31AA031189 (ZT). The authors wish to thank members of the Miles laboratory for their helpful advice and assistance during the course of this work.

## Supplementary Figures and Tables are available from the authors upon request

**Suppl. Figure S1:** Block-colored dendrogram of TOM at deep-splits 0-4.

**Suppl. Figure S2:** Main Gene Ontology categories, paired by semantic similarity, represented by genes significantly differentially expressed at FDR ≤ 0.1 between High-Drinkers and Low-Drinkers.

**Suppl. Figure S3:** Main Gene Ontology categories, paired by semantic similarity, represented by genes significantly differentially expressed at FDR ≤ 0.1 between Low-Drinkers and Control.

**Suppl. Figure S4: a)** Scale-free fit of log-scaled TPM values **b)** Scale-free fit of gene counts after variance stabilizing transformation (VST)

**Suppl. Figure S5:** Heatmap of overlap analysis between Zeisel single-cell RNAseq dataset and WGCNA modules.

**Suppl. Figure S6:** Manhattan plot of chromosomal locations of all cis and trans eQTLs. Location based on the location of the genetic marker, points colored by location of the gene.

**Suppl. Figure S7: a)** Chromosomal location of confidence intervals of QTLs in Chr 18, trans band located between 30 Mb and 90 Mb and percentage of QTLs with location included among their CI (trans band 1). **b)** Chromosomal location of confidence intervals of QTLs in Chr 18, trans band located between 20 Mb and 70 Mb and percentage of QTLs with location included among their CI (trans band 2). **c)** Chromosomal location of confidence intervals of QTLs in Chr 13, trans band located between 50 Mb and 120 Mb and percentage of QTLs with location included among their CI. **d)** Chromosomal location of confidence intervals of QTLs in Chr 2, trans band located between 40 Mb and 120 Mb and percentage of QTLs with location included among their CI.

**Suppl. Figure S8:** REVIGO treemap plot of Gene Ontology enrichment of trans band 1 on Chr 18.

**Suppl. Figure S9:** REVIGO treemap plot of Gene Ontology enrichment of trans band 2 on Chr 18.

**Suppl. Figure S10:** REVIGO treemap plot of Gene Ontology enrichment of trans band on Chr 13.

**Suppl. Figure S11:** Plot of LOD scores for significant eigengene QTLs on Chr 4, Chr 5, and Chr 13 with haplotype analysis plots.

