## Supplementary material for "Identification of Genetic and Genomic Influences on Progressive Ethanol Consumption in Diversity Outbred Mice": DO_Mice_SupplFigS8.pdf

Chr 18 trans band 1, GO: Biological Process

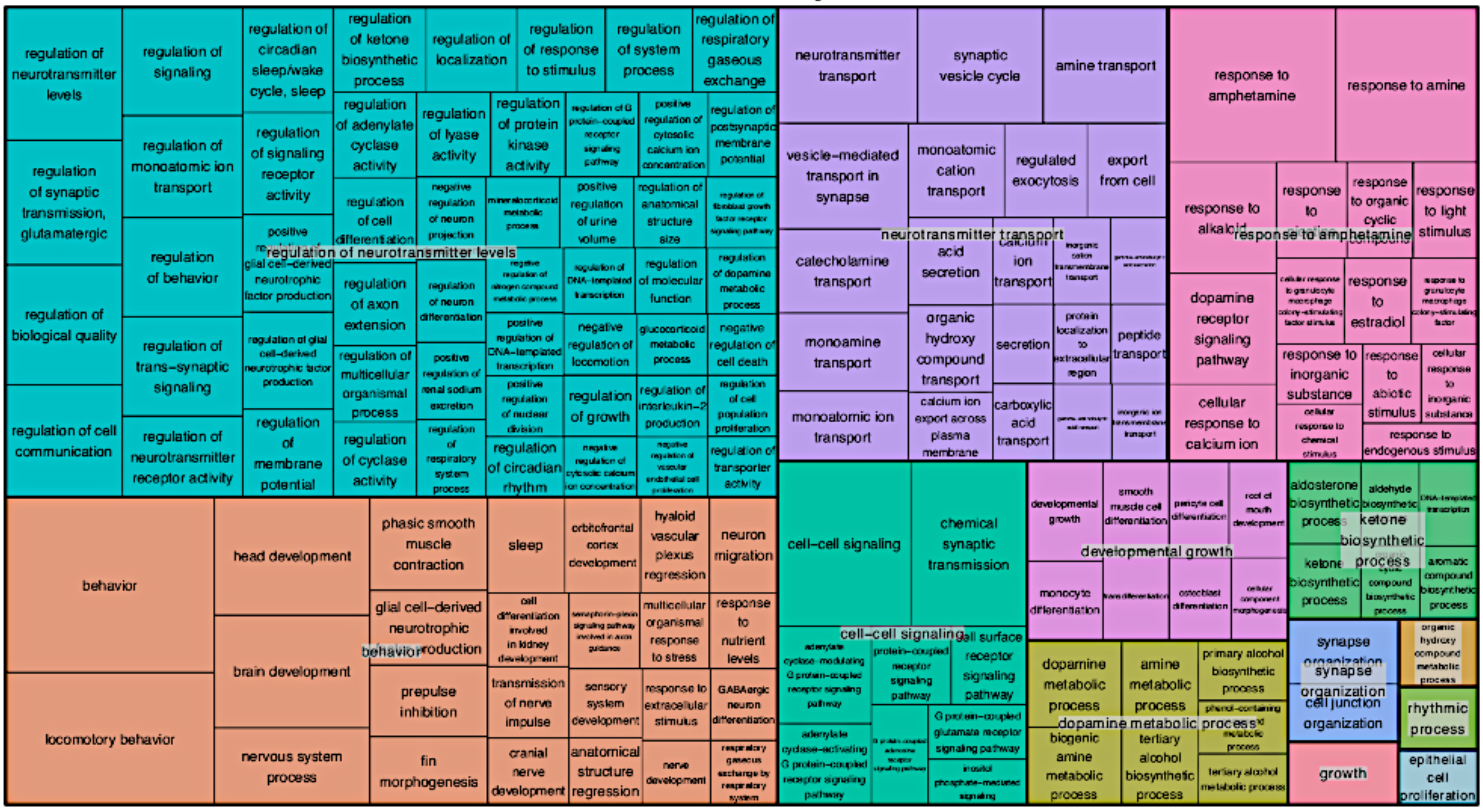


Chr 18 trans band 1, GO: Cellular Component

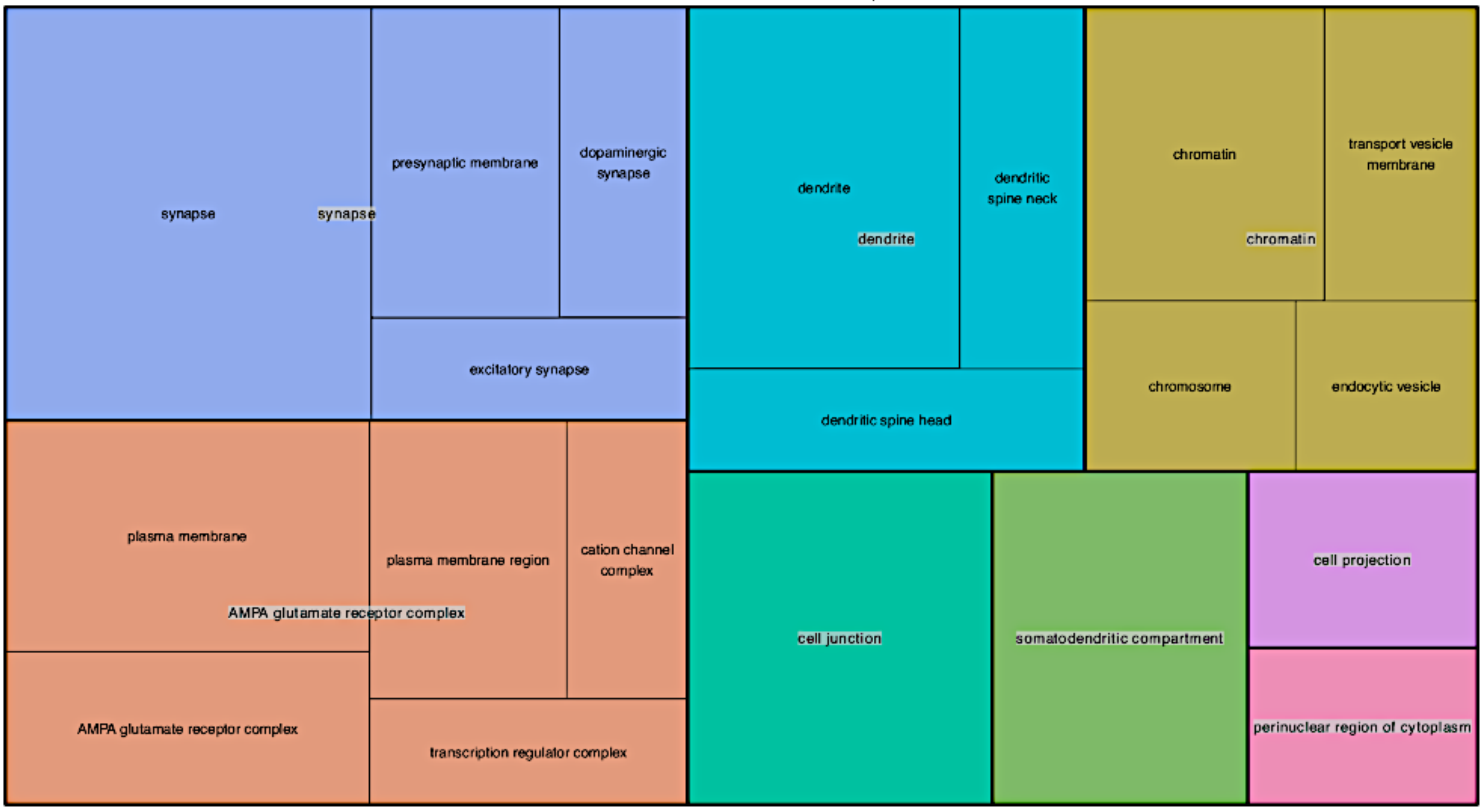


Chr 18 trans band 1, GO: Molecular Function

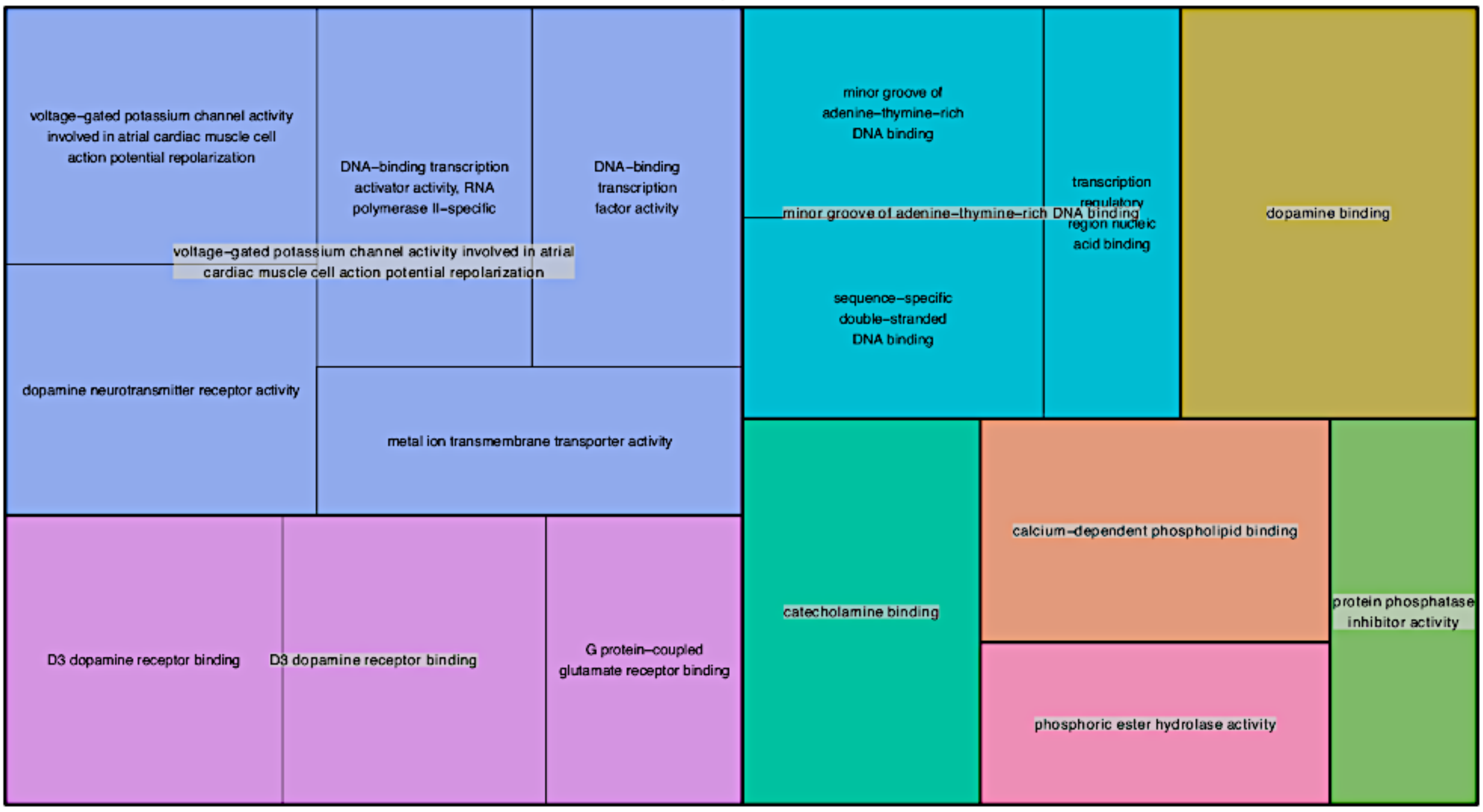
