## Supplementary figures and images for "Identification of Genetic and Genomic Influences on Progressive Ethanol Consumption in Diversity Outbred Mice"

### DO_Mice_SupplFigS1.pdf

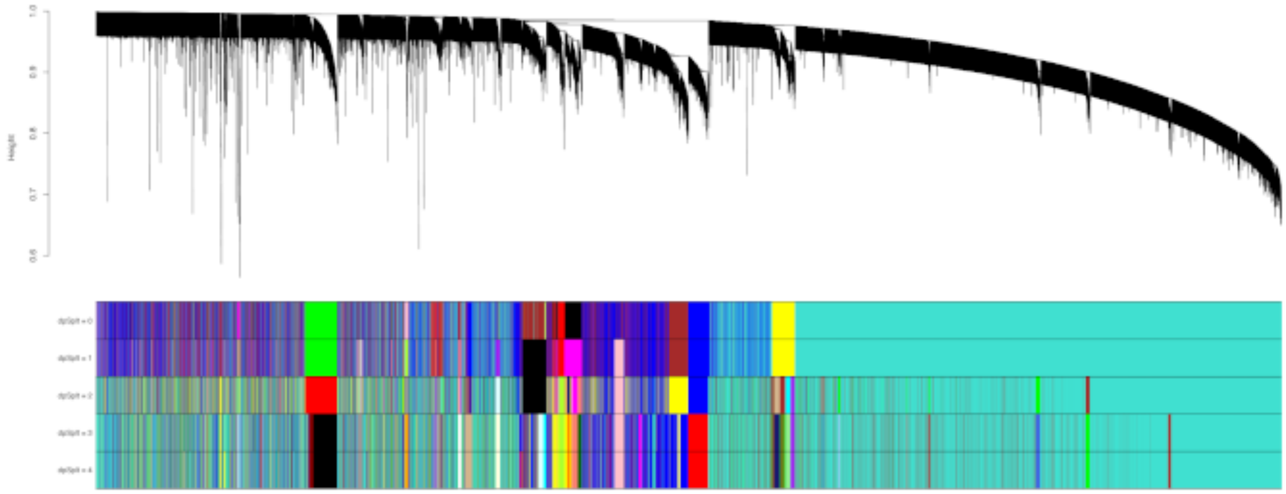

MDS plot DS3

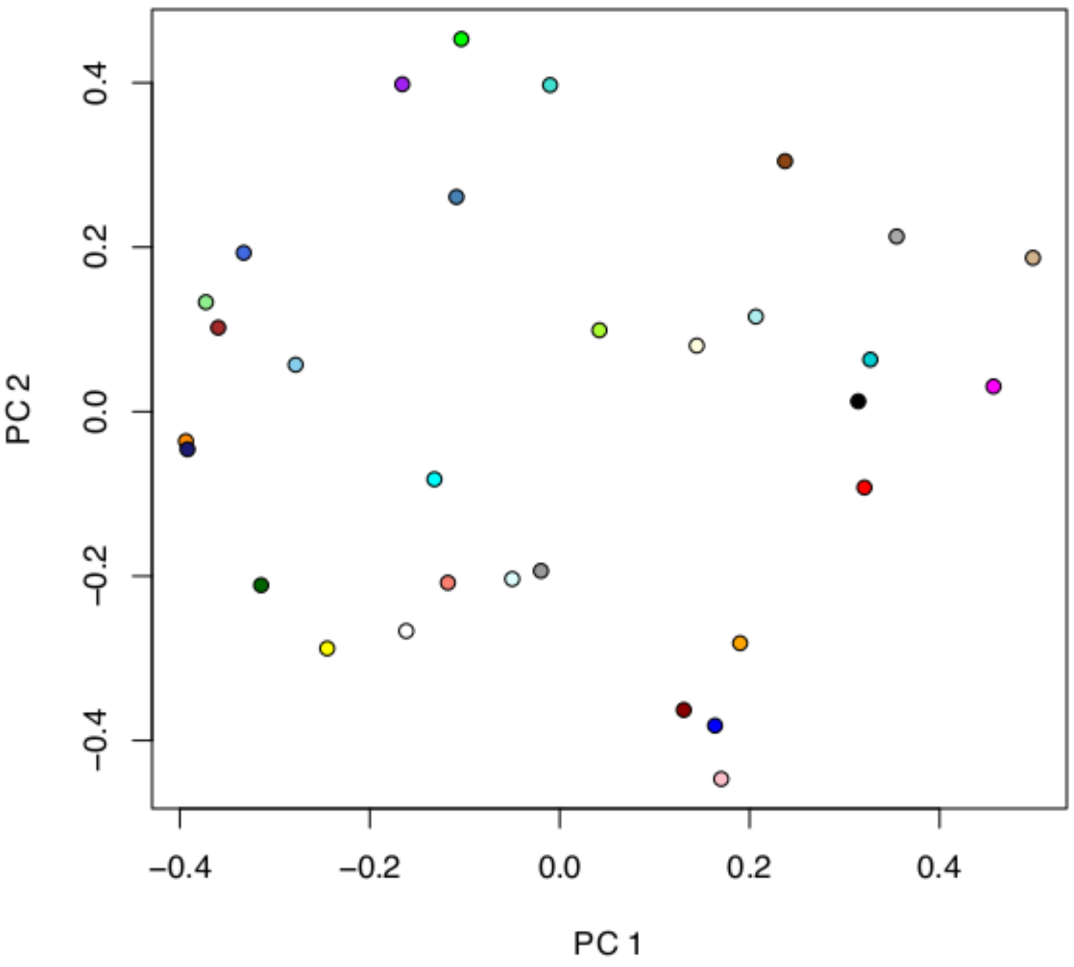

### DO_Mice_SupplFigS2.pdf

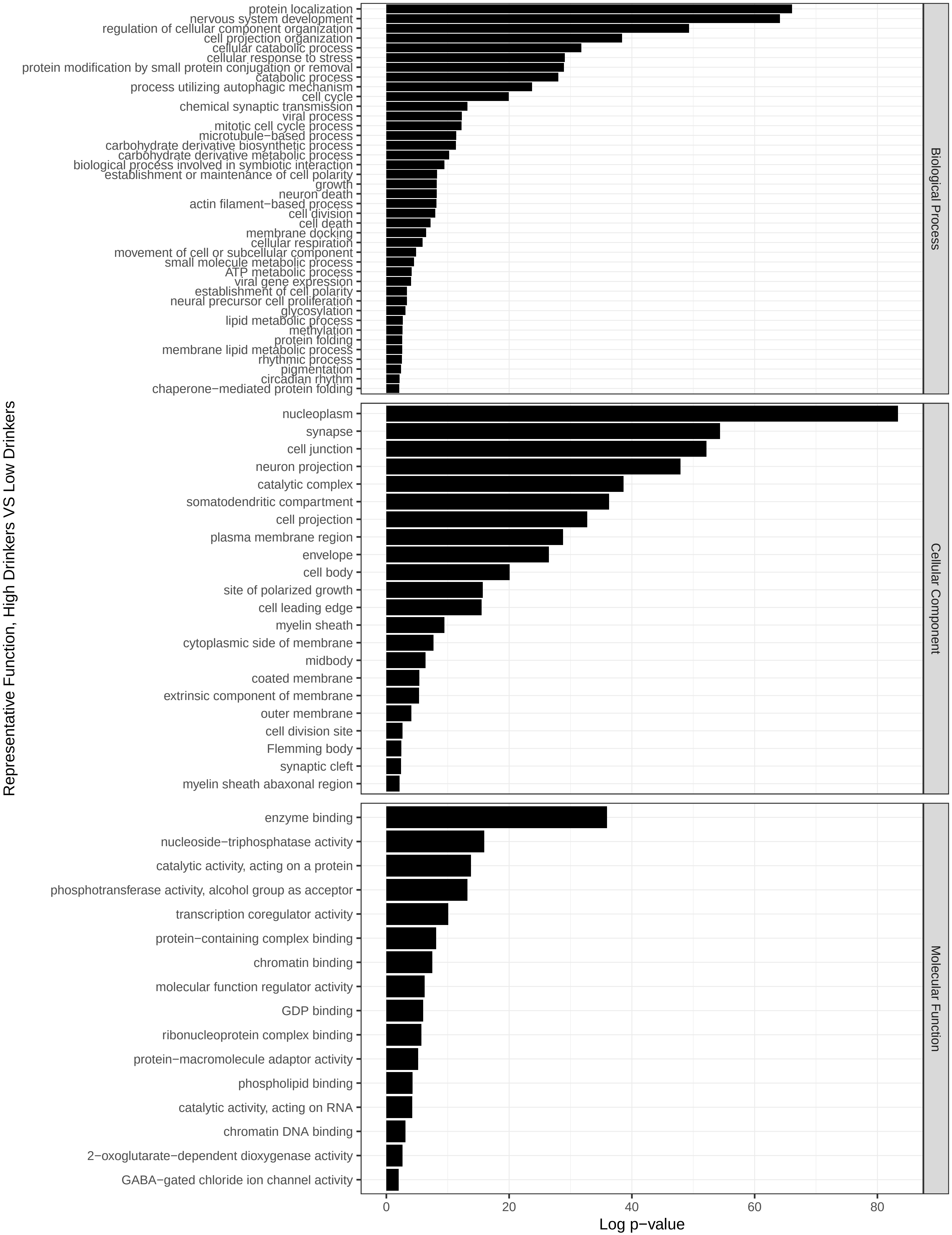

### DO_Mice_SupplFigS3.pdf

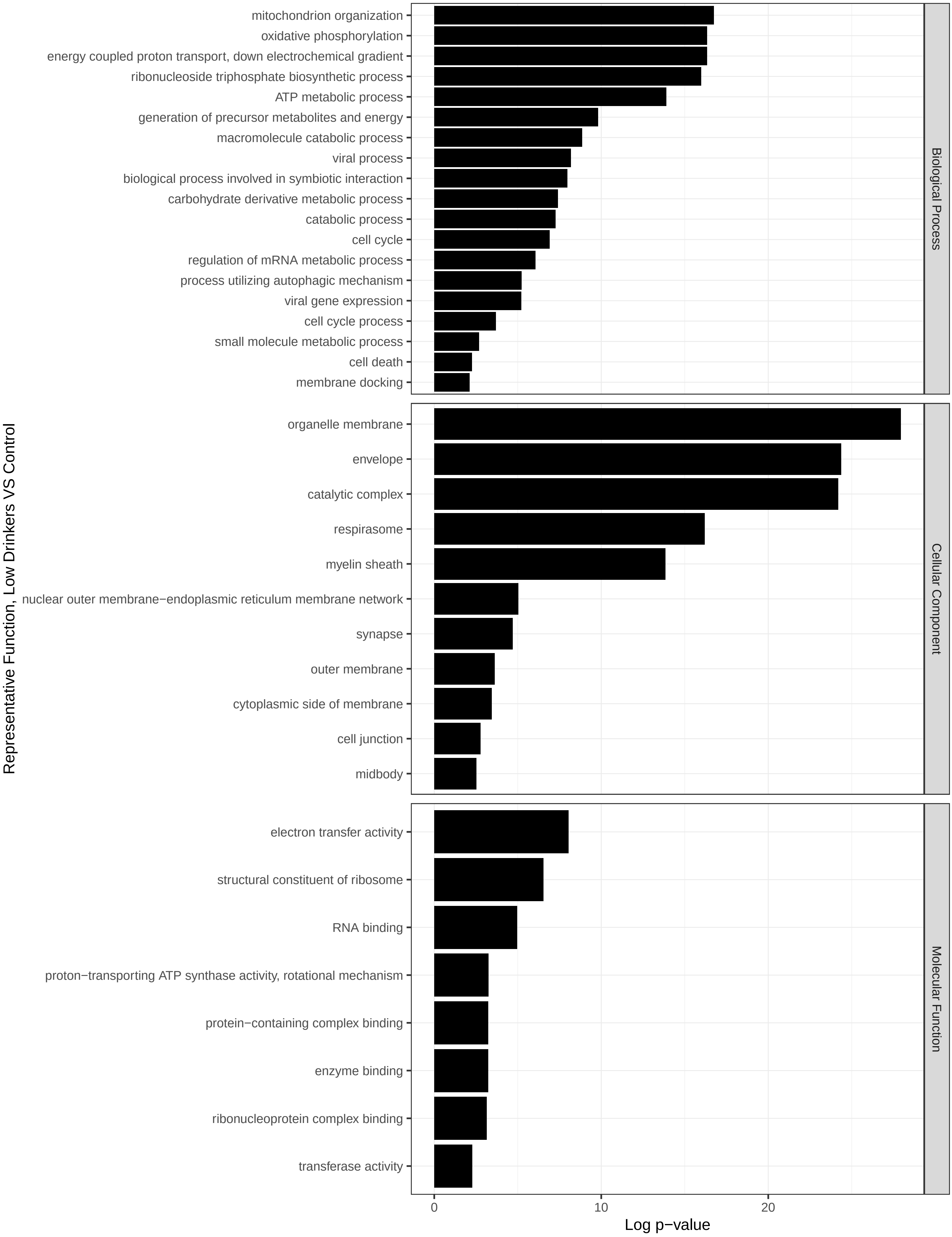

### DO_Mice_SupplFigS4.pdf

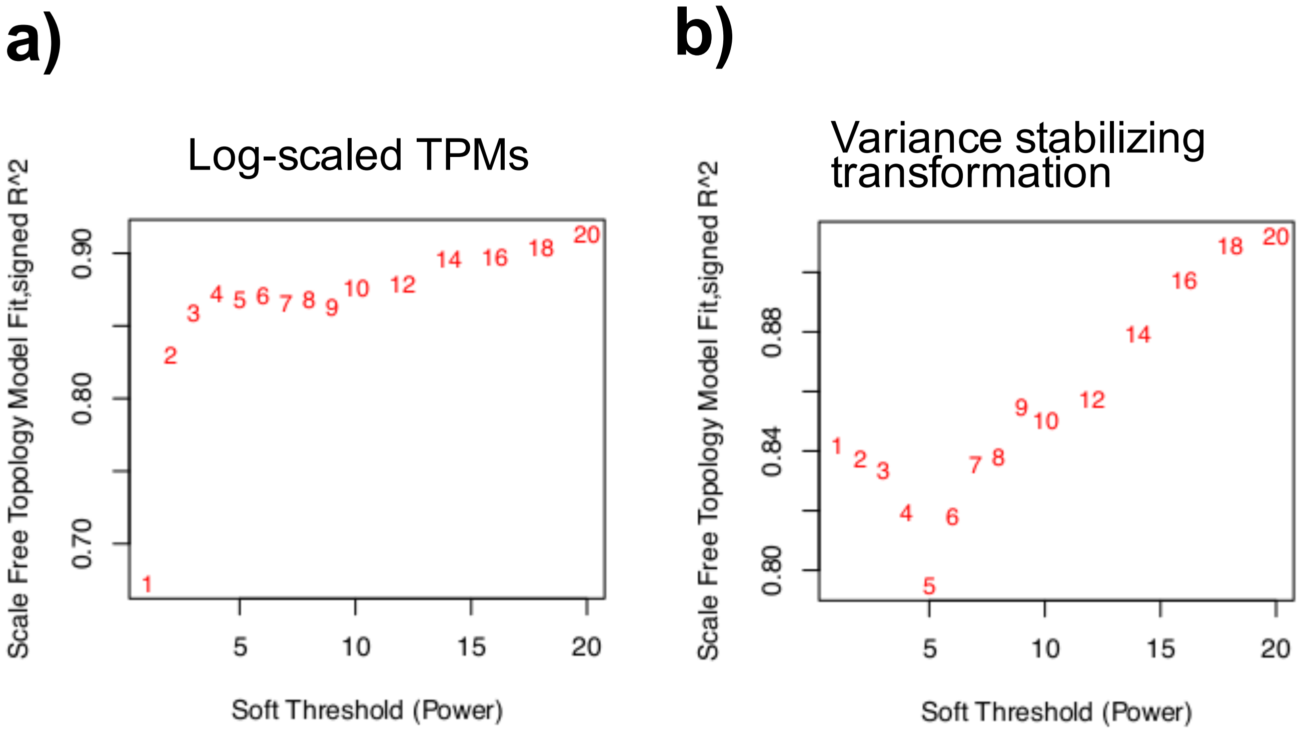

### DO_Mice_SupplFigS5.pdf

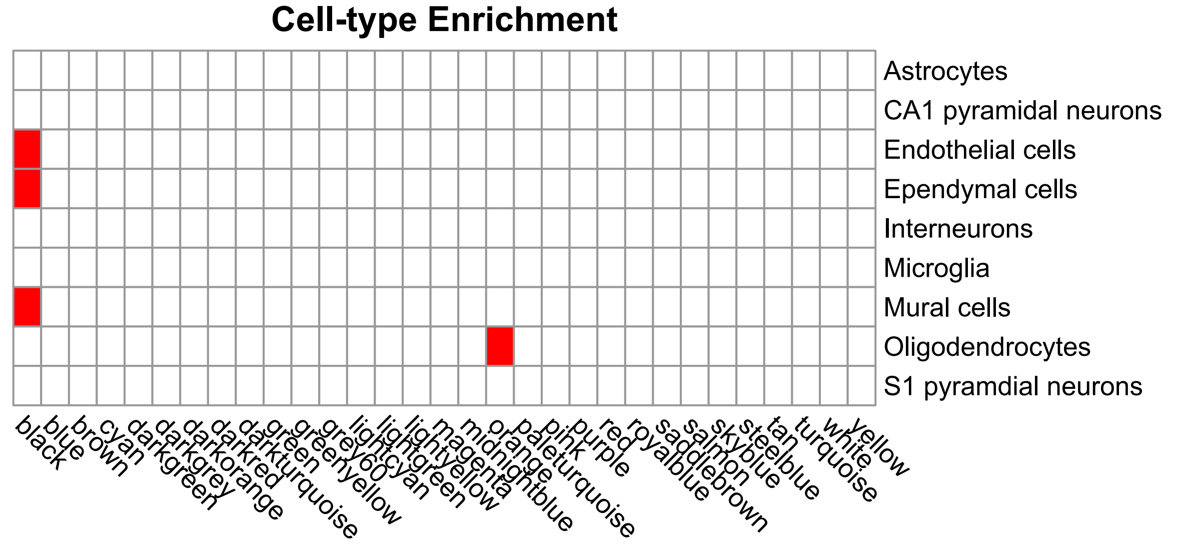

### DO_Mice_SupplFigS6.pdf

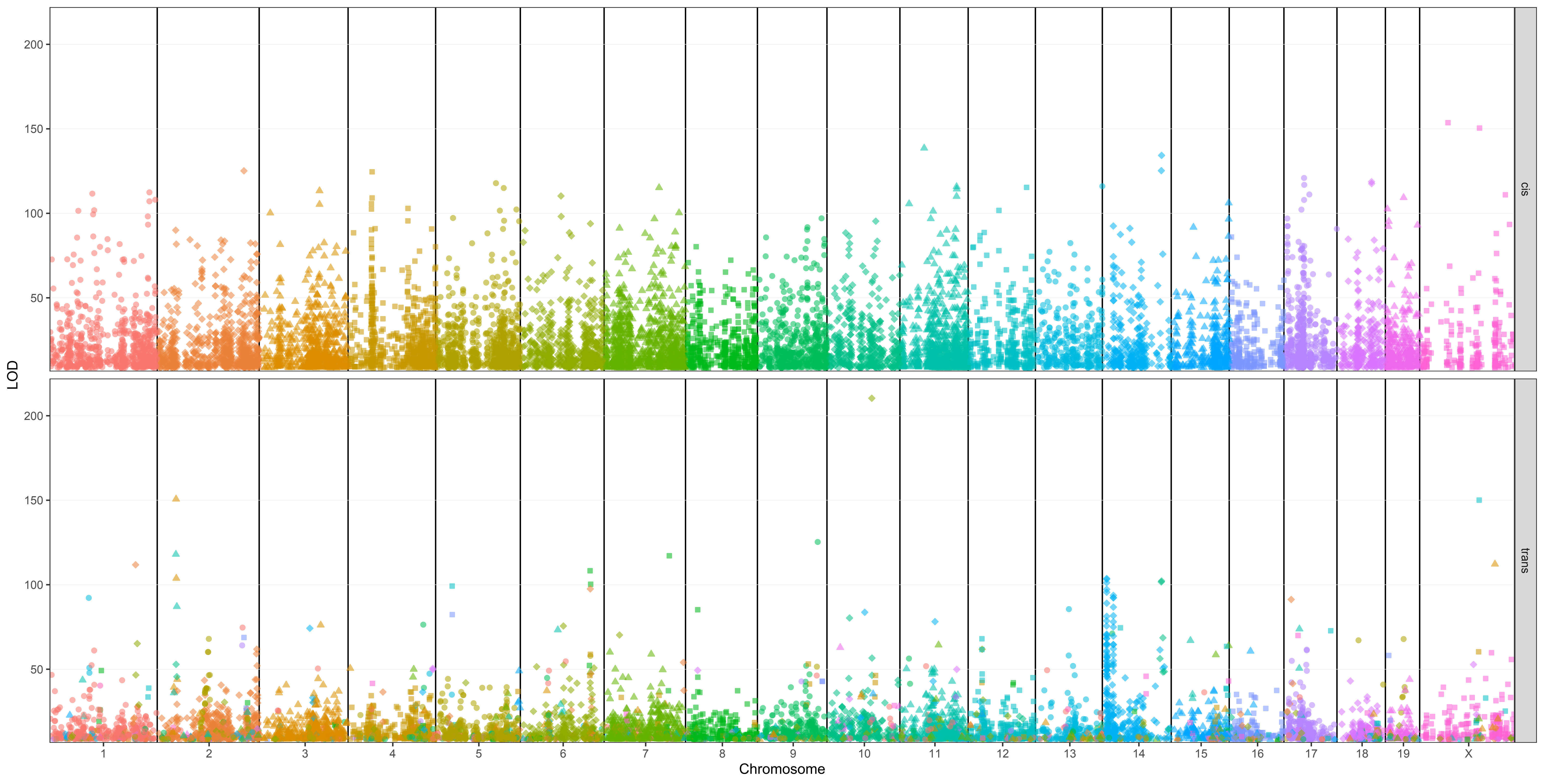

### DO_Mice_SupplFigS7.pdf

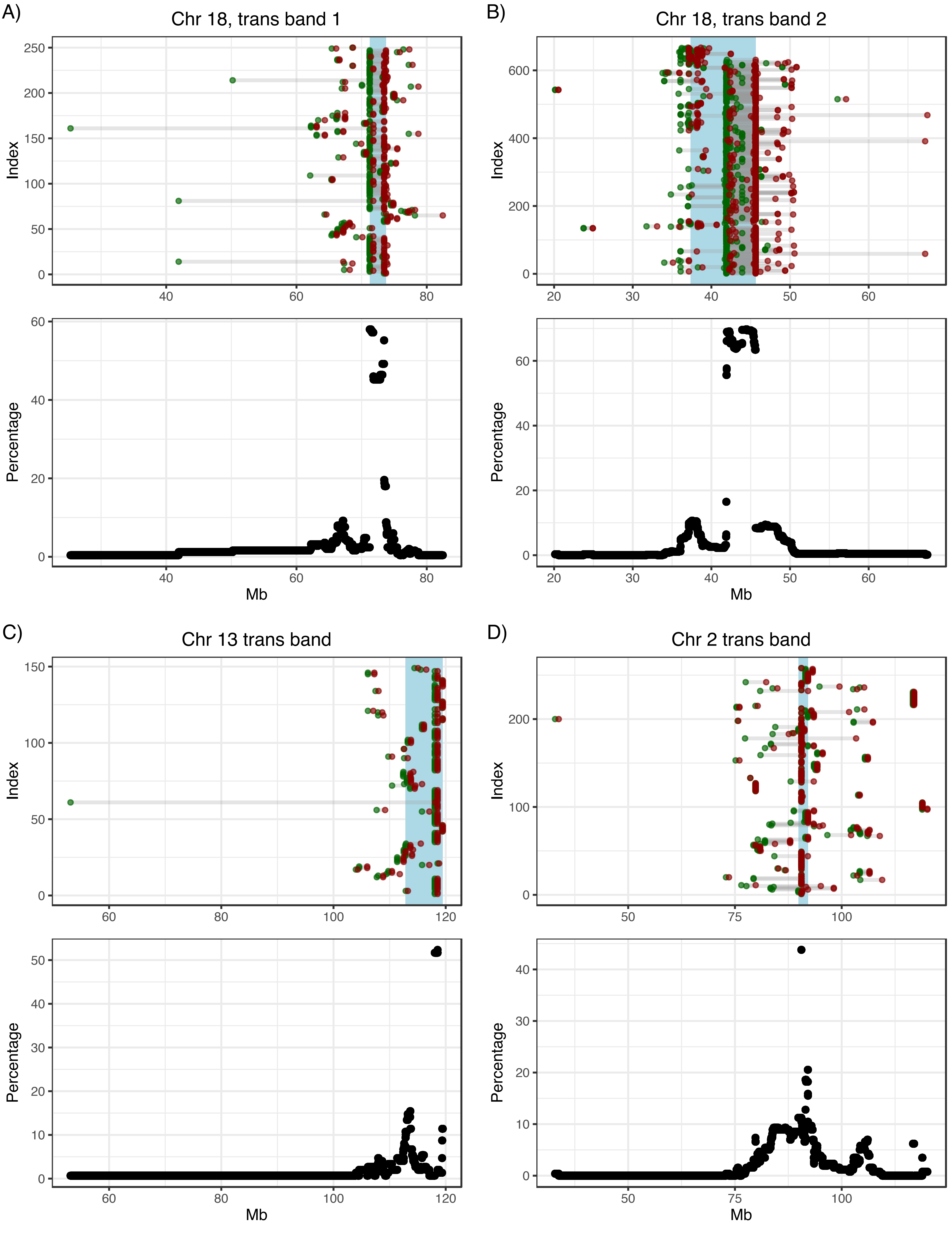

### DO_Mice_SupplFigS10.pdf

Chr 13 trans band, GO: Biological Process

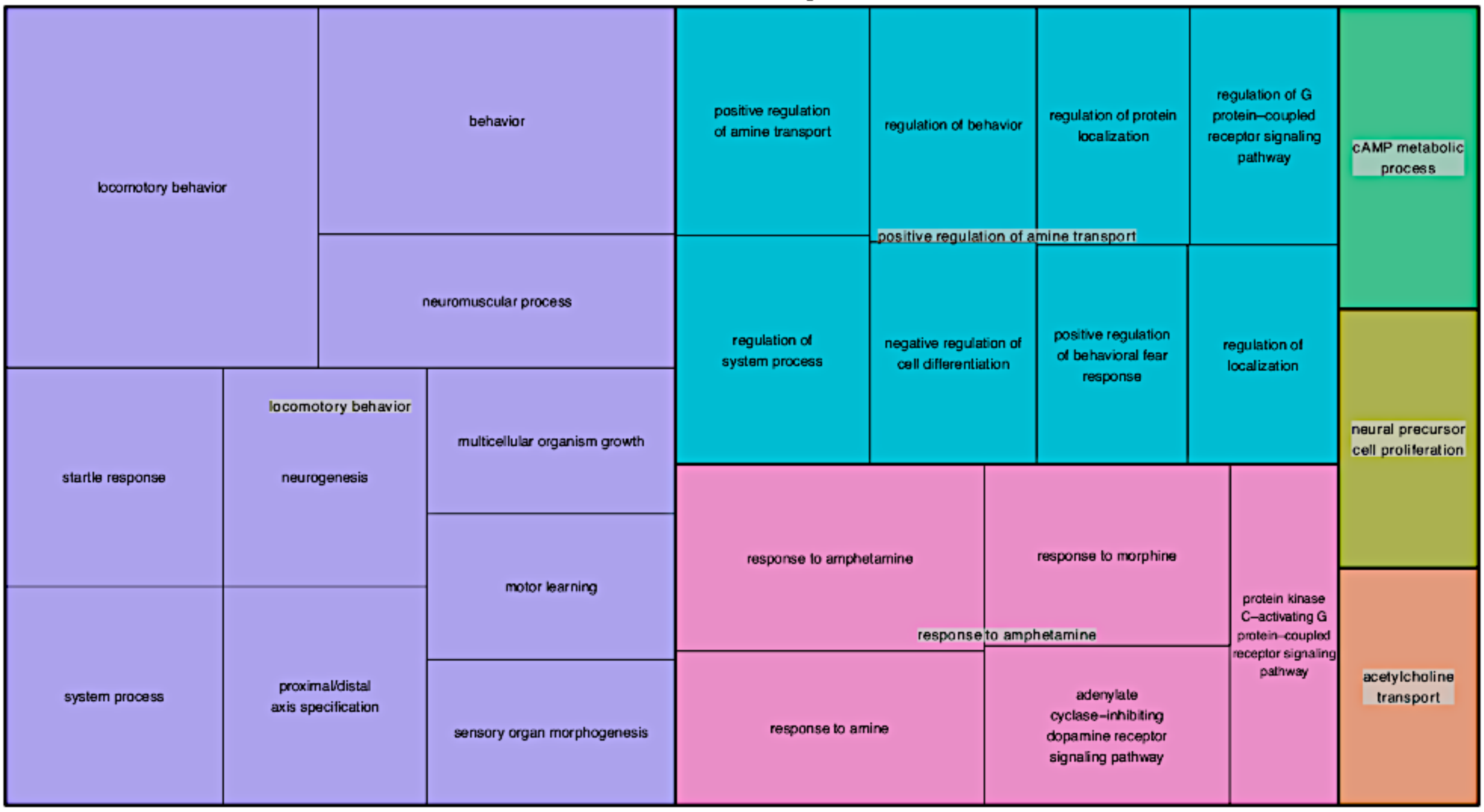


Chr 13 trans band, GO: Molecular Function

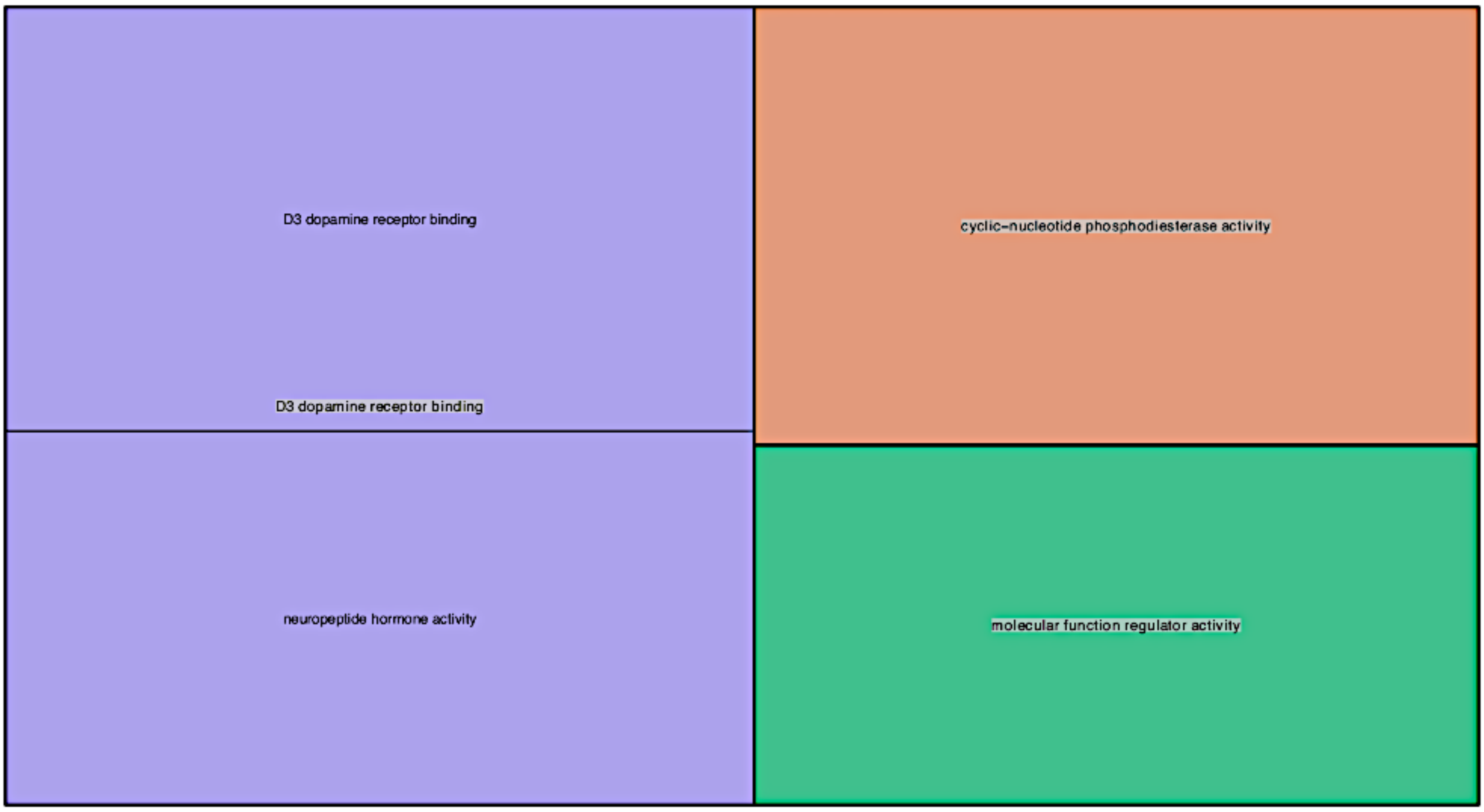

### DO_Mice_SupplFigS11.pdf

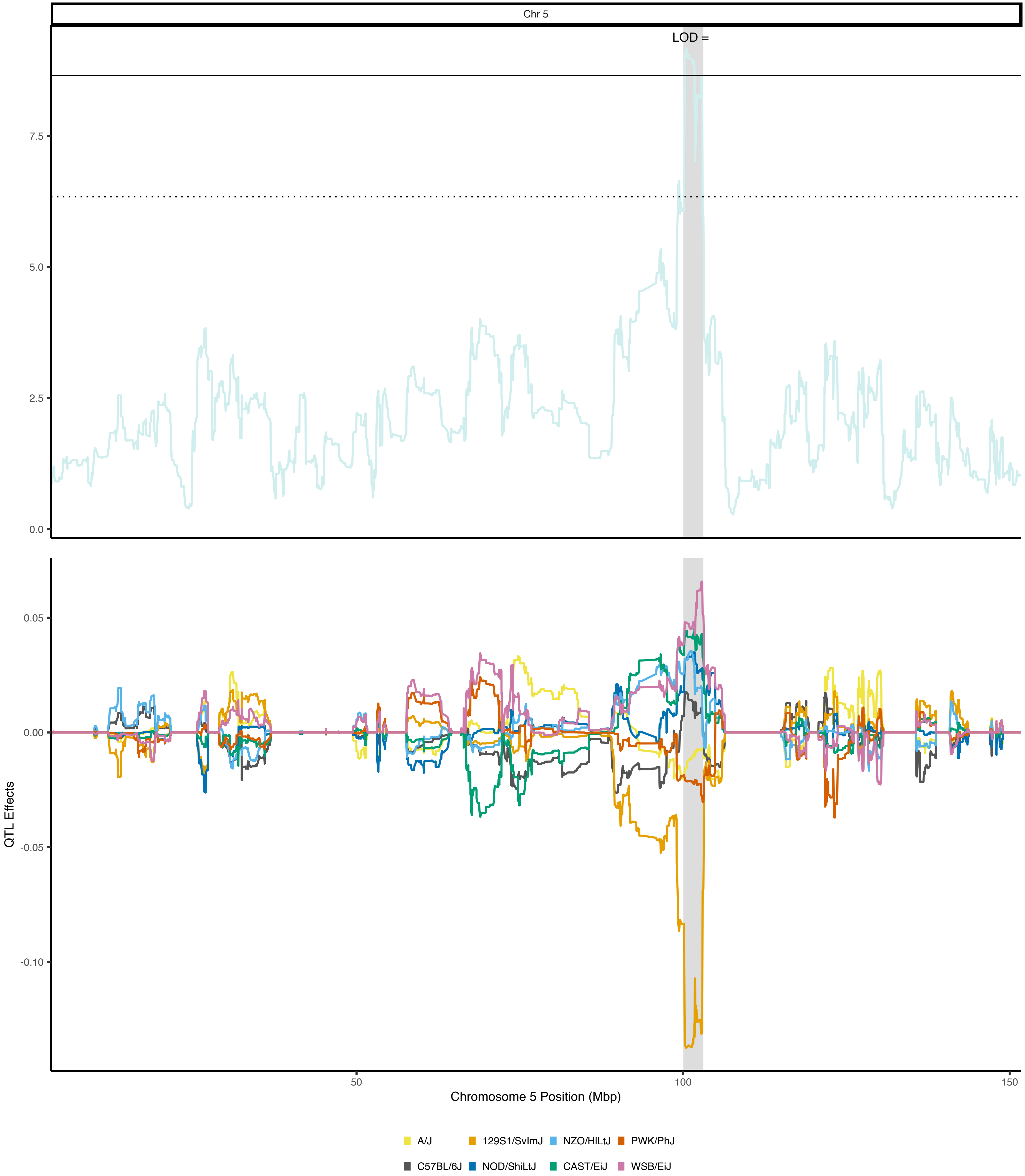

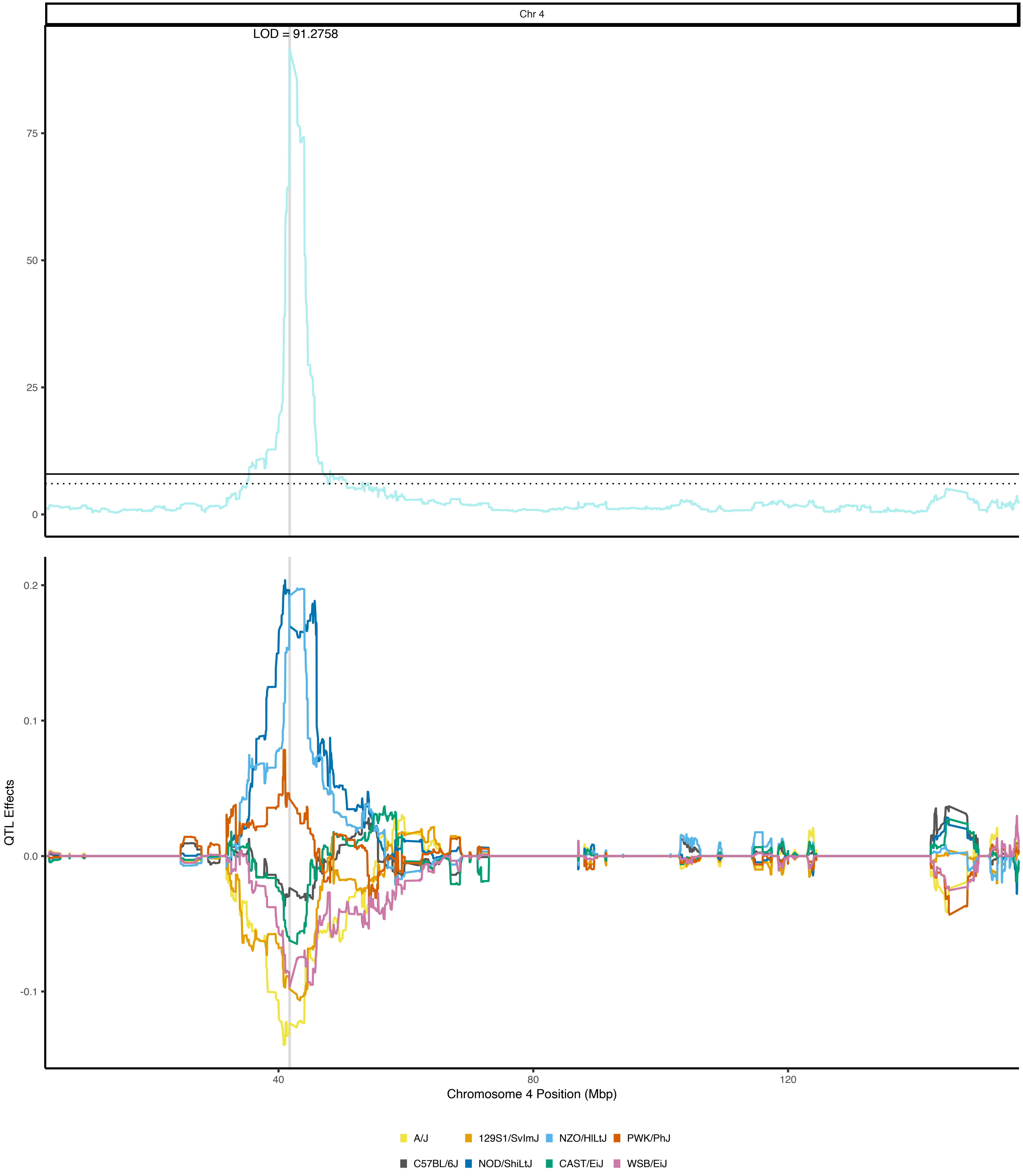

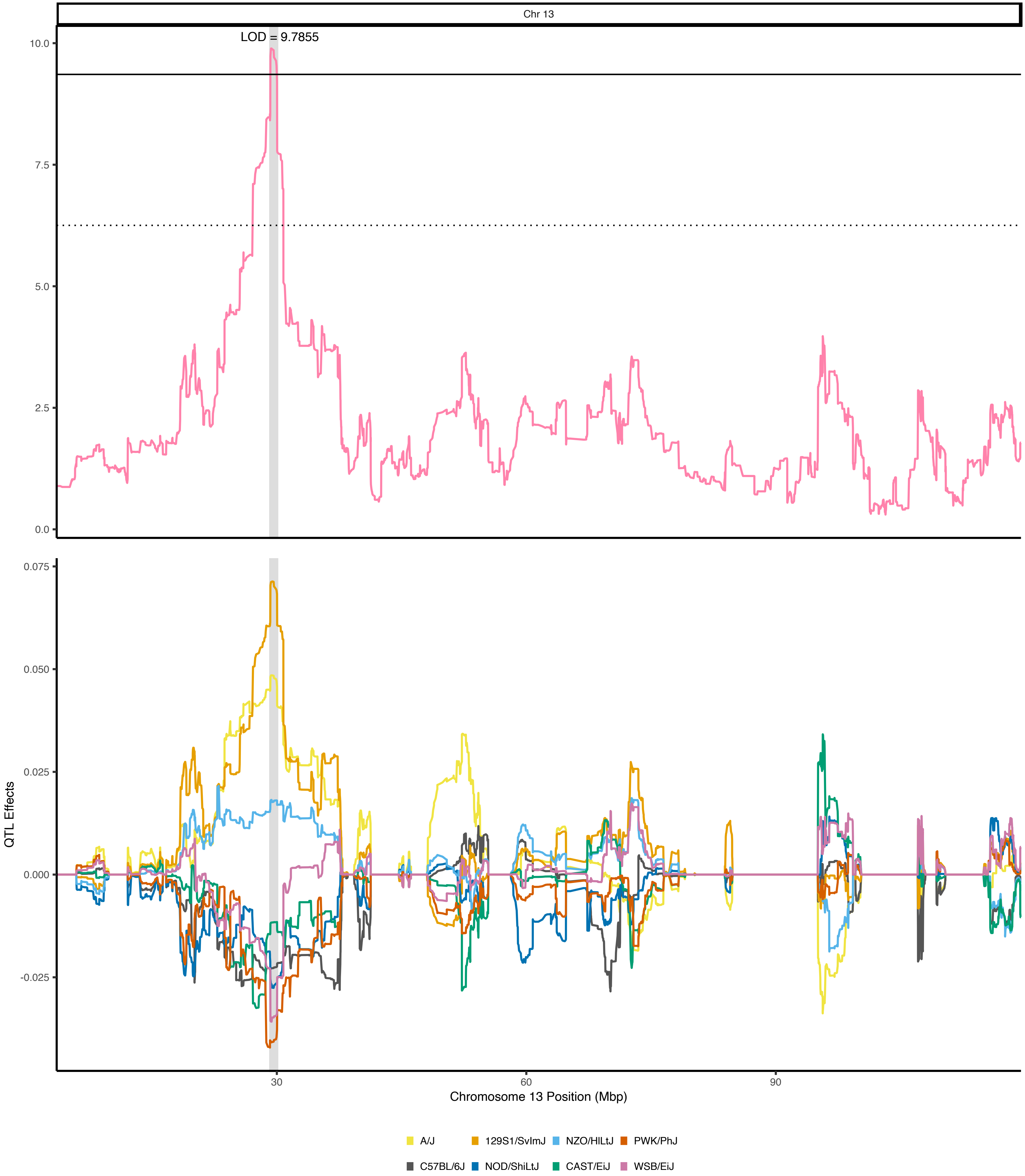
